# The statistical structure of the hippocampal code for space as a function of time, context, and value

**DOI:** 10.1101/615203

**Authors:** Jae Sung Lee, John Briguglio, Sandro Romani, Albert K. Lee

## Abstract

Hippocampal activity represents many behaviorally important variables, including context, an animal’s location within a given environmental context, time, and reward. Here we used longitudinal calcium imaging in mice, multiple large virtual environments, and differing reward contingencies to derive a unified probabilistic model of hippocampal CA1 representations centered on a single feature – the field propensity. Each cell’s propensity governs how many place fields it has per unit space, predicts its reward-related activity, and is preserved across distinct environments and over months. The propensity is broadly distributed—with many low, and some very high, propensity cells —and thus strongly shapes hippocampal representations. The result is a range of spatial codes, from sparse to dense. Propensity varied ~10-fold between adjacent cells in a salt-and-pepper fashion, indicating substantial functional differences within a presumed cell type. The stability of each cell’s propensity across conditions suggests this fundamental property has anatomical, transcriptional, and/or developmental origins.

## Introduction

How the brain represents the world is one of the central questions of neuroscience. Representations in many primary sensory areas are characterized by localized receptive fields organized in a topographic manner (Penfield and Boldrey, 1937; Talbot and Marshall, 1941; Mountcastle, 1957; Hubel and Wiesel, 1962). In the hippocampus, an animal’s location in a given environment (context) is represented by place cells and their spatial receptive fields (place fields) (O’Keefe and Dostrovsky, 1971). Unlike primary visual or somatosensory areas, however, hippocampal place cell representations do not have a topographic organization (e.g. nearby cells have fields that are far apart) (O’Keefe, 1976). Furthermore, individual place cells can have multiple fields in an environment (O’Keefe, 1976; Fenton et al., 2008; Rich et al., 2014), fields of multi-field cells are randomly dispersed across the environment (Rich et al., 2014), and cells may have no fields (i.e. are “silent”) in some environments (O’Keefe and Conway, 1978; Thompson and Best, 1989; Wilson and McNaughton, 1993; Epsztein et al., 2011; Alme et al., 2014; Rich et al., 2014). Therefore, hippocampal representations possess several features that appear random – something that is shared by association areas and other regions (Rigotti et al., 2013).

What are the organizing principles of such distributed representations? On the one hand, randomness itself has value. For instance, if place fields of adjacent cells were always near each other, this would greatly reduce the number of possible representations of environmental contexts, whereas random representations maximize memory capacity and minimize interference between representations. On the other hand, hippocampal representations exhibit clearly nonrandom structure. This structure, in contrast to the geometric organization of primary sensory areas, only becomes apparent at the statistical level due to the randomness in the number of fields and their locations. In particular, the propensity of a hippocampal CA1 neuron to have a place field in each unit of space varies greatly across cells and follows a simple, highly skewed gamma distribution (Figure 1A), such that (i) many cells have a low propensity, some have high propensity, others have in-between values, and (ii) given its propensity, a cell’s fields are distributed randomly in space (i.e. a spatial Poisson process) (Rich et al., 2014). Similar distributions have been found in other hippocampal regions (Alme et al., 2014; Witharana et al., 2016) as well as the olfactory system (Turner et al., 2008), suggesting such skewed distributions may be a general property (Buzsaki and Mizuseki, 2014) of representations across many brain areas (where, for nonspatial areas, the propensity would indicate the probability that a neuron participates in the representation for each item of concern).

**Figure 1.**
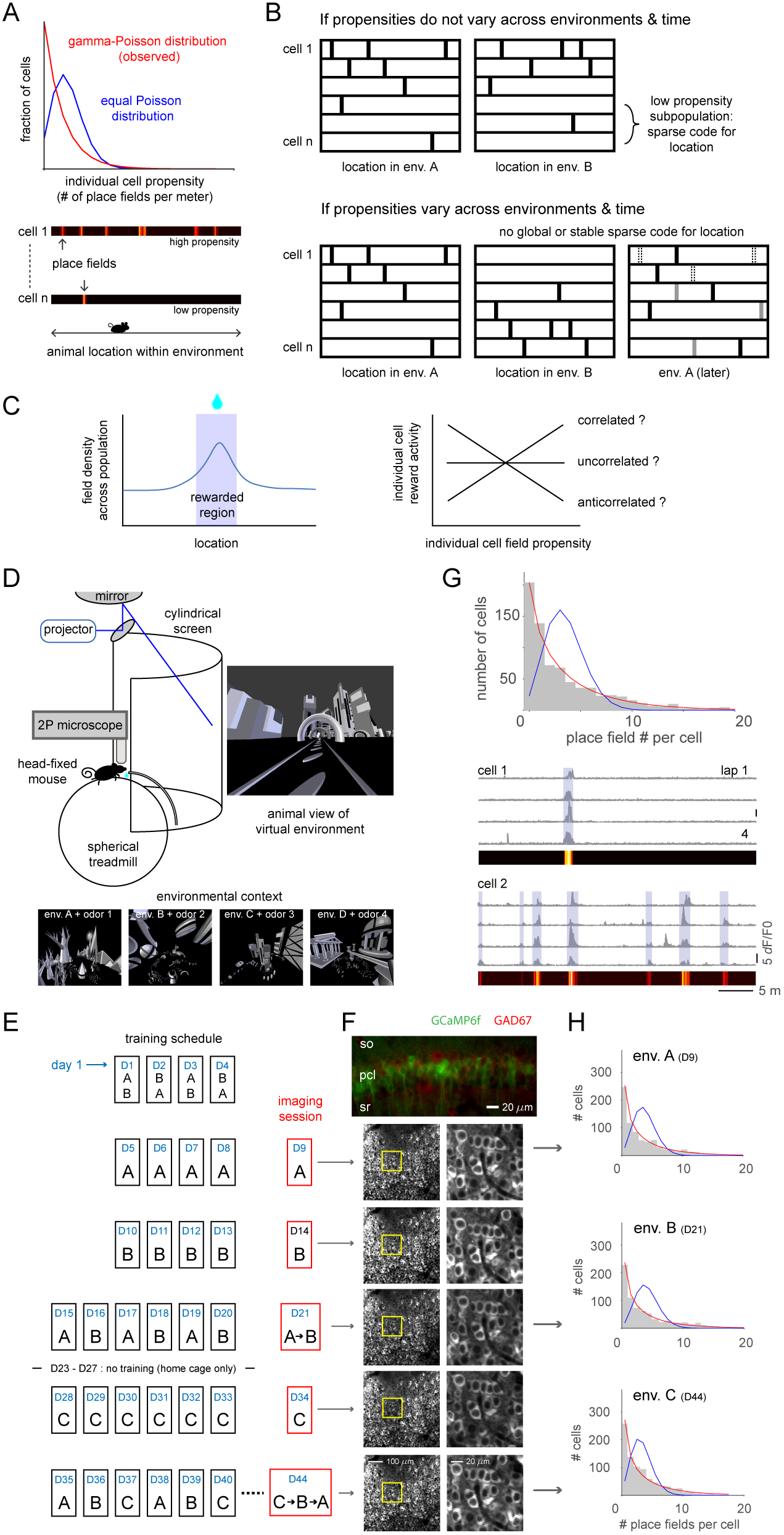
Skewed distribution of place field propensity of individual dorsal hippocampal CA1 pyramidal cells in multiple large virtual environmental contexts observed with longitudinal two-photon calcium imaging. (A) Number of place fields per cell assuming cells have same (equal Poisson, blue) or gamma-distributed Poisson (observed, red) rate of forming place fields per unit length. (B) Schematic showing preservation of each cell’s propensity across environments and time would yield sparse code for location (black: place fields; gray: new fields; open boxes: fields that disappeared). (C) Schematic of known increase in field density in consistently rewarded locations (left) and possible relationship between propensity and reward activity. (D) Schematic of experimental apparatus (top) and large virtual environments (bottom). (E) Training and imaging schedule in 3 different environments from day 1 (D1) to 44 (D44). (F) GCaMP6f (green) and GAD67 (red) signal (top) and standard deviation images of GCaMP6f traces in lap 1 (left column). Imaging session indicated by arrows from (E). Expanded images of yellow boxes (right column). (G) Place field number distribution in an environment (top). Fit of equal-Poisson (blue) or gamma-Poisson (red) distribution. Example place cell with 1 field (middle) and cell with 7 fields (bottom). (H) Place field number distribution in 3 different environments on different days. See also Figures S1-S3.

One potential function of a highly skewed distribution of field propensity is that the many low propensity cells result in a sparse code for location (e.g. a substantial fraction of cells have exactly one place field across a large extent of space) (Rich et al., 2014), and sparse coding has been shown to have desirable properties (Tsodyks and Feigelman, 1988; Olshausen and Field, 2004; Katkov et al., 2017). However, for sparse coding to work in this way, the same set of cells would need to have low field propensity across different environments as well as over time (Figure 1B). More generally, if each cell’s propensity were preserved across different conditions, this would indicate a significant degree of structure underlying all hippocampal representations.

Separately, hippocampal representations have been shown to be influenced by sensory and value cues. CA1 place field density is elevated in locations where reward is consistently provided (Hollup et al., 2001; Dupret et al., 2010; Danielson et al., 2016; Zaremba et al., 2017; Gauthier and Tank, 2018; Sato et al., 2018), which could support the hippocampus’ role in navigating to spatial goals (Morris et al., 1982). Previous work on field propensity used environments in which value was spatially homogeneous (Rich et al., 2014; Alme et al., 2014; Witharana et al., 2016). How do value-based modulations in overall field density relate to each cell’s field propensity (Figure 1C)?

Here we bring together these many features that characterize hippocampal CA1 representations to develop a unified statistical description of place field structure with respect to time, context, and value. We employed 2-photon calcium imaging of the hippocampus in awake head-fixed mice (Dombeck et al., 2010; Lovett-Barron et al., 2014; Sheffield and Dombeck, 2015; Basu et al., 2016; Danielson et al., 2016, Malvache et al., 2016; Gauthier and Tank, 2018; Hainmueller and Bartos, 2018; Sato et al., 2018) as they navigated multiple large virtual environments. Large environments (Kjelstrup et al., 2008) allowed high-resolution measurement of field statistics such as propensity (Fenton et al., 2008; Rich et al., 2014) and virtual reality enabled use of many arbitrarily large environmental contexts. Furthermore, imaging allowed propensities of large numbers of individual cells to be tracked for months (Ziv et al., 2013; Rubin et al., 2015). Finally, by manipulating reward contingencies in these environments, we investigated the interaction of value and propensity. The result of our analysis is a simple probabilistic model centered on the field propensity that can predict the structure of hippocampal representations under a wide variety of conditions.

## Results

### Long-term calcium imaging in multiple large virtual environments

Mice were trained to run on a 40 cm-diameter spherical treadmill at the center of a half-cylindrical screen onto which monochromatic virtual environments were projected (Figure 1D) (Harvey et al., 2009; Dombeck et al., 2010; Cohen et al., 2017). The animal’s view of the environment was rendered with custom virtual reality software (Cohen et al., 2017) and updated based on treadmill movement. We designed 4 different environments with distinctly shaped ~40 m-long tracks surrounded by unique objects (Figures 1D and S1). Each animal explored 3 of the environments (A, B, C, where A represents the first environment explored regardless of its identity; Figure 1E), with each environment accompanied by a distinct background odor cue. Behavior consisted of running the entire track’s length in one direction then being teleported to the start for another lap, for a total of 4-8 laps in a given environment per session. Animals explored each environment over multiple days/sessions. All analyses were performed on sessions in which the environment was familiar (≥4 days of previous experience), unless otherwise noted. In the first set of experiments, water reward was provided at random locations that changed for each lap. We imaged the calcium activity of dorsal hippocampal CA1 pyramidal neurons in transgenic mice (n = 4) expressing GCaMP6f (Figure 1F; top: GCaMP6f-expressing cells do not express GAD67) (Dana et al., 2014). 700-870 neurons were identified in each imaging session and we could follow 125-430 neurons across all sessions for up to 3 months (Figure 1F).

### Field propensity varies greatly across individual cells

For the first element of the model, we quantified the distribution of the number of place fields per neuron. As reported previously (Rich et al., 2014), the distribution of field numbers across the population was highly skewed, well-fit with a gamma-Poisson distribution, and far different and far more spread out than predicted by a (equal-)Poisson distribution (Figures 1G and S2), which is what would be expected if each cell had the same propensity to have a place field per unit space. Therefore, different cells have clearly different field propensities. Furthermore, given its propensity, each cell’s fields were distributed randomly in space (spatial Poisson per cell, Figure S2). These results using calcium imaging in head-fixed mice exploring familiar virtual environments thus match those seen in previous extracellular recordings from freely moving rats running on a novel (i.e. previously unencountered) 48 m track (Rich et al., 2014). In addition, the skewed field number distribution was not changed by time or environmental context (Figures 1H and S2, K-S test, p > 0.15 for all comparisons) and not explained by the moderate change in mean field number along proximal-distal (Henriksen et al., 2010) or superficial-deep (Mizuseki et al., 2011) axes (Figure S3).

**Figure 2.**
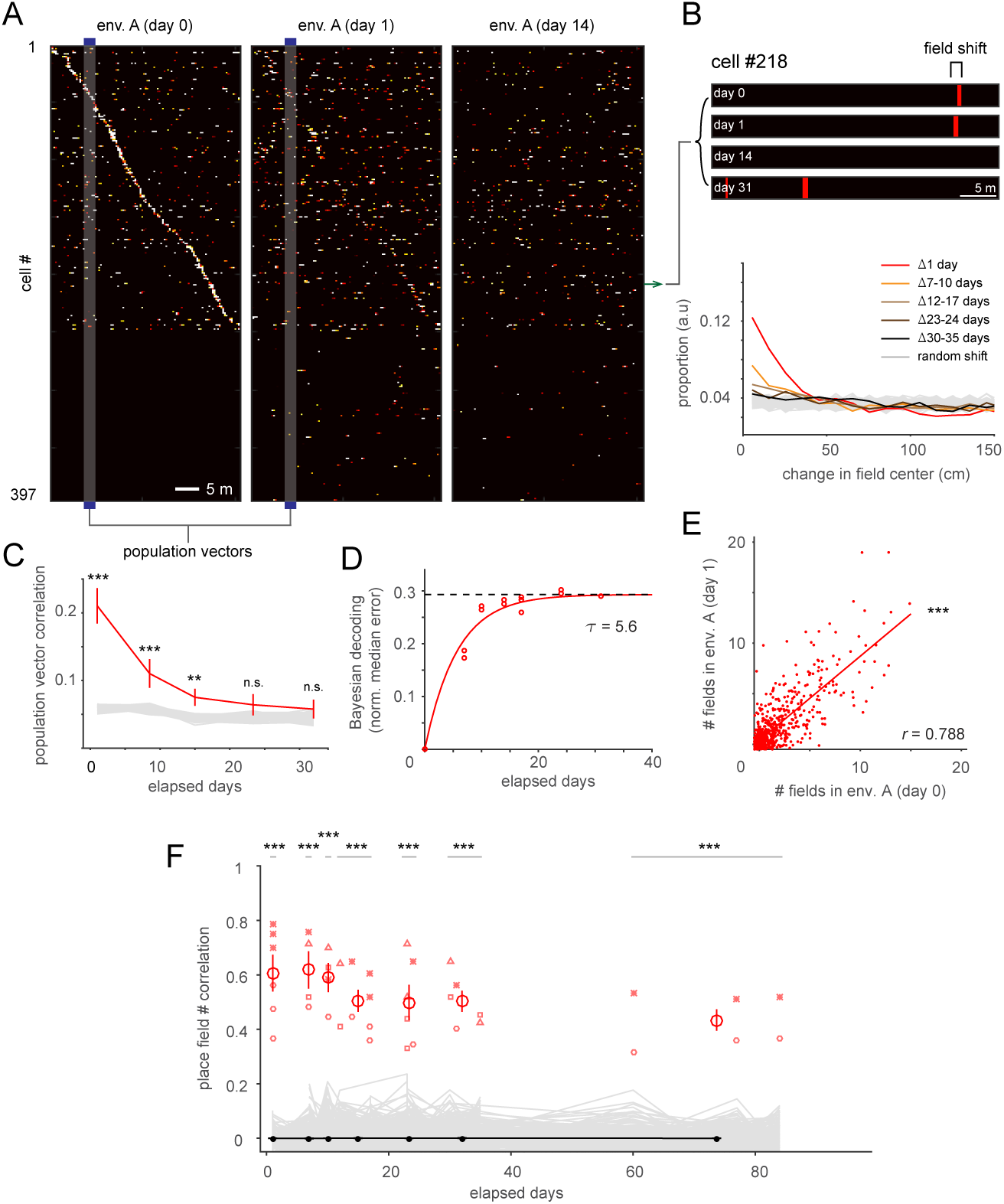
Field propensity of each cell is preserved over time in the same environmental context, even while the place field representation of that environment gradually evolves. (A) Each cell’s calcium activity in laps 1-4 were averaged on each day. Cells were ordered by location of peak activity on day 0. Same cell order was kept for days 1 and 14. (B) Example cell field shift in same environment over time (top). Shift of center of fields pooled across all pairs of sessions and animals (n = 23417-38063 field pairs for each group of days, bottom). (C) Average correlation between binary place field population vectors for same location in same environment on different days (mean ± s.e.m., n = 4 mice; gray: shuffled data). (D) Normalized median error of Bayesian decoding of animal location on one day based on calcium activity from a different day. (E) Scatterplot where each point displays one cell’s place field numbers on 2 different days in same environment. (F) Correlation of number of fields per cell across pairs of sessions in same environment (e.g. correlation of scatterplot in (E); small red marks: individual session pairs, unique symbol per mouse; large red circles and error bars: mean ± s.e.m.; gray: shuffled data; black circles and error bars: shuffled mean ± s.e.m.). *p < 0.05, **p < 0.01, ***p < 0.001.

### Individual cells preserve their field propensity over time

Since different cells had widely different propensities, we asked if each cell’s propensity was preserved in a given environment over time. First, we checked the stability of place field locations in these large environments. As seen in freely moving animals exploring small (~1 m) environments (Mankin et al., 2012; Ziv et al., 2013; Mankin et al., 2015; Rubin et al., 2015; Hainmueller and Bartos, 2018), CA1 fields reliably represented the same location for ~2 weeks but also gradually changed over time. This phenomenon was quantified in several different ways – the location of the center of each place field (Figures 2A and 2B), the correlation between population vectors (PV) of the simultaneous activity pattern across all cells at corresponding locations (Figure 2C), and Bayesian decoding of location (Figure 2D) (Davidson et al., 2009) (statistical significance here and elsewhere is reported in figures, e.g. using asterisks, if not in the main text).

While individual field locations changed such that decoding performance gradually decreased to chance levels after ~2 weeks, surprisingly the field propensity of each cell was preserved across months in a given environmental context. Specifically, the (Pearson) correlation coefficient of the number of place fields of each cell (Figure 2E) was significantly higher than chance for at least 2 months (Figure 2F, 1 day, n = 6, p < 0.001; 60-84 days, n = 6, p < 0.001).

### Individual cells preserve their field propensity across distinct environmental contexts

We next tested whether each cell’s field propensity was preserved across different environmental contexts. Beforehand, we checked that place field representations in different large environments were distinct (global remapping) (O’Keefe and Conway., 1978; Muller and Kubie., 1987; Leutgeb et al., 2005), as in standard-sized (~2-4 m-long) virtual environments (Danielson et al., 2016; Cohen et al., 2017, Gauthier and Tank, 2018, Hainmueller and Bartos, 2018). We compared representations after switching between familiar environments during a session (Figure 3A). In sharp contrast to the similarity of place field locations in the same environment on different days (Figure 2), representations of different environments were uncorrelated as assessed by the same measures (field center locations, PV correlation, Bayesian decoding) (Figures 3B-3D).

**Figure 3.**
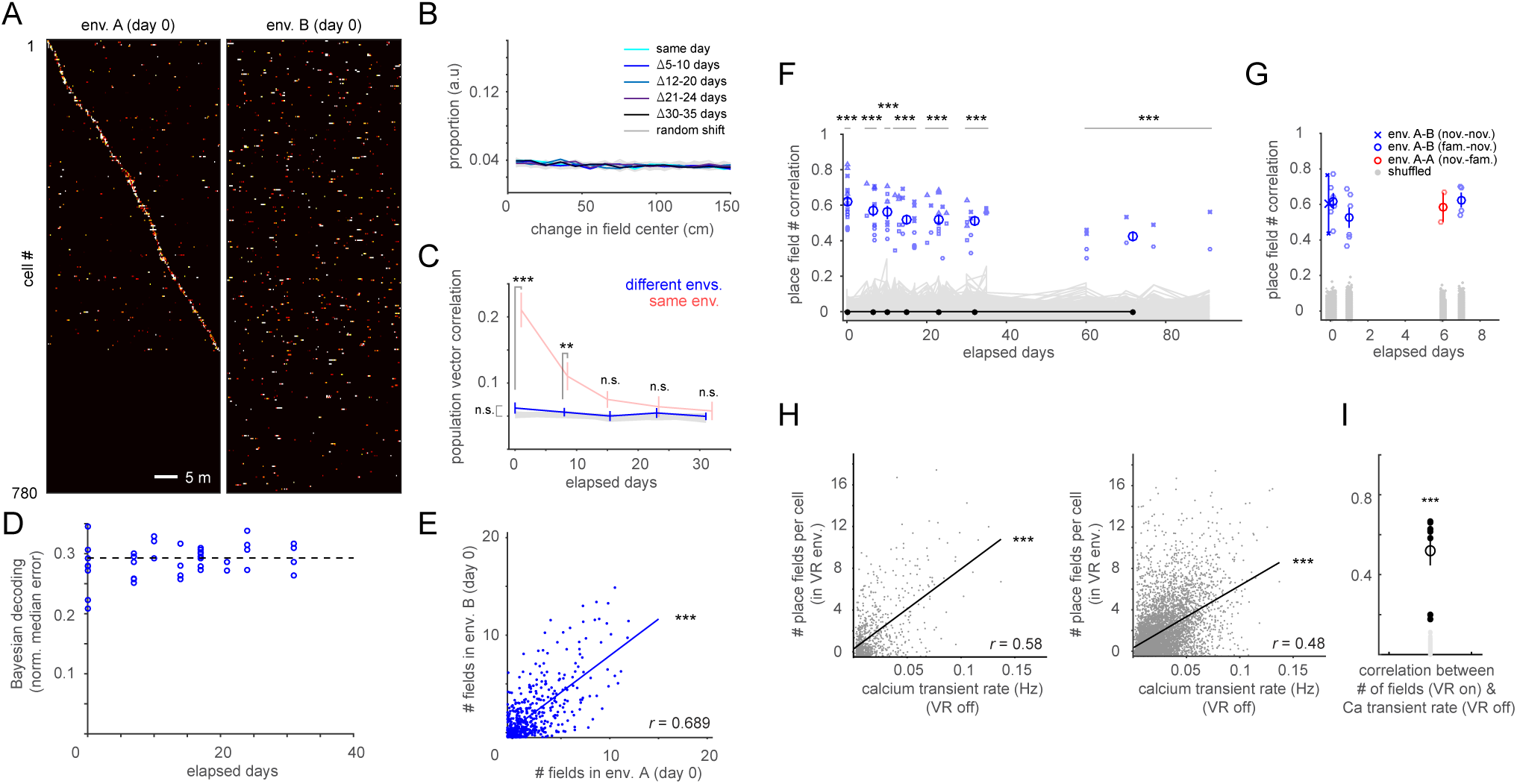
Field propensity of each cell is preserved across distinct environmental contexts, even though the place field representations are completely different. (A)-(F) Similar to Figures 2A-2F except for different environmental contexts. (G) Place field number correlation between different environments where one or both are novel (mean ± s.e.m.). (H) Scatterplot of calcium transient rate in dark and subsequent virtual exploration session field number per cell (example session from a mouse, left; pooled over 8 sessions, 2 each from 4 mice, right). (I) Correlation as in (H), left, for each session (n = 8 sessions, 2 each from 4 mice; black points: individual sessions; black open circle with error bar: mean ± s.e.m.; gray points: shuffle). *p < 0.05, **p < 0.01, ***p < 0.001.

While ensemble place codes were completely distinct for different large environments, each cell had similar numbers of place fields in all contexts (Figures 3E and 3F). This is surprising as fields have been shown to be controlled by sensory inputs (O’Keefe and Conway., 1978; Muller and Kubie., 1987; Deshmukh and Knierim, 2013; Geiller et al., 2017) and the visual features of each environment were very different. The similarity in field numbers is consistent with extracellular recordings showing correlations in average firing rates between two environments explored on the same day (Mizuseki and Buzsaki, 2013). Furthermore, by tracking cells across time with imaging, we could determine that field propensity was preserved across contexts for as long as we performed the experiments (up to 90 days, Figure 3F). The preservation of propensity also applied to distinct environments explored for the first time (Figure 3G).

As propensity was preserved, we could combine field numbers of each cell across multiple environments and sessions to give the highest resolution measure of propensity. The resulting propensities across the population in a single animal ranged by over an order of magnitude (gamma fit 90:10 percentile propensity ratio range: 6.1-48.0, mean: 24.4; versus Poisson fit range: 1.7-2.2, mean: 1.9, n = 4 mice) (Figures S3B-S3D).

For a subset of sessions, we placed the animal on the treadmill but in the dark and with no odor cue before turning on the virtual environment. The activity level of each cell was correlated with its field propensity (Figures 3H and 3I), suggesting preservation of propensity was not dependent on any visual features that may have been similar between the otherwise visually different environments.

Overall, the time course of the field number correlation was similar for same and different environments (Figure 4A). Furthermore, the average calcium transient rate of each cell in a session, irrespective of where in the environments the individual transients occurred, was highly correlated across sessions widely separated in time and followed similar time courses whether in the same or different environments (Figure 4B). Together, these results suggest that the overall excitability of each CA1 neuron is the determining factor for place field propensity.

**Figure 4.**
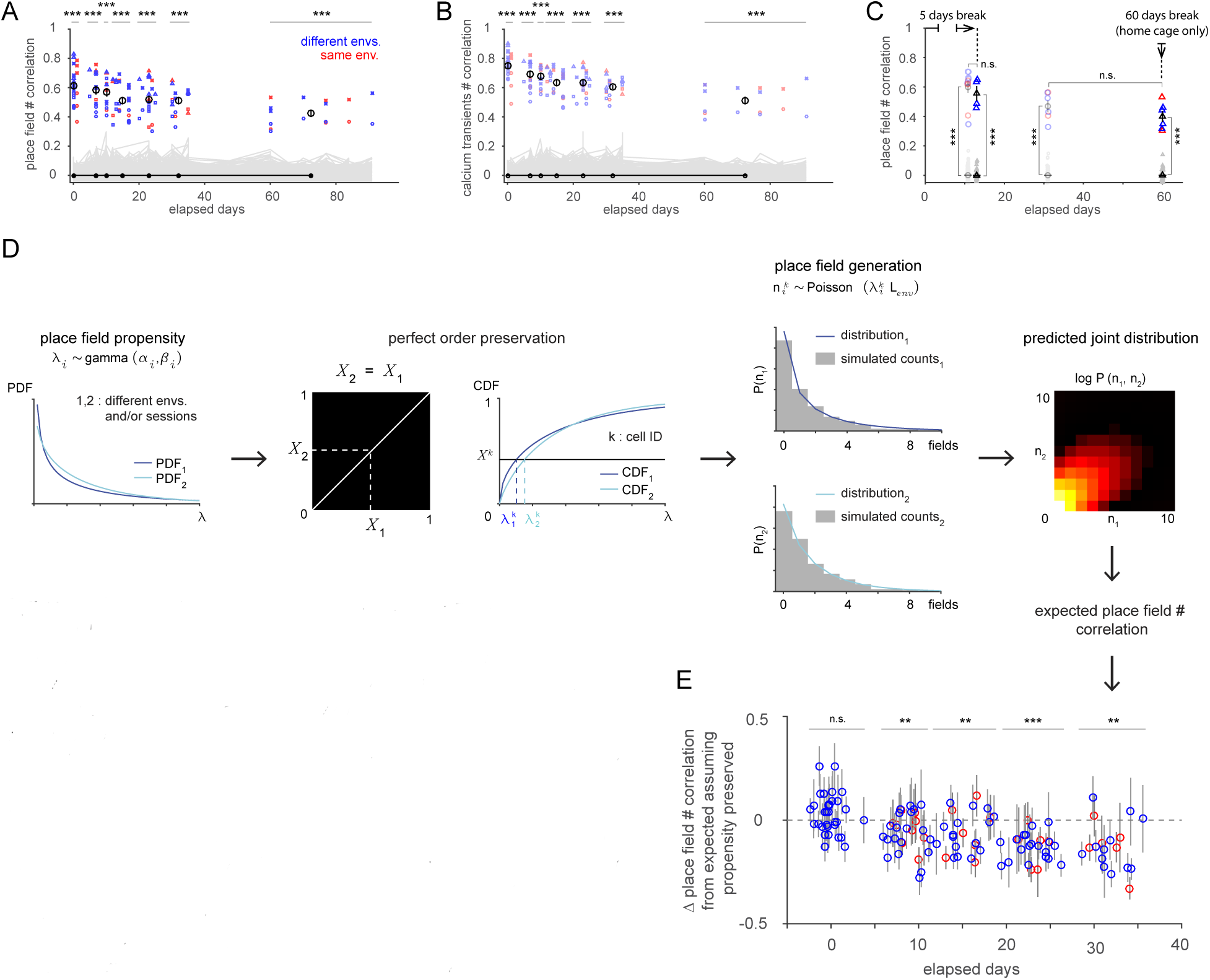
Propensity of each cell is highly preserved across days and environments. (A) Place field number correlation in same (Figure 2F, red) or different (Figure 3F, blue) environments superimposed. (B) Same as (A) except calcium transient number correlation irrespective of locations where transients occur. (C) Place field number correlation between sessions with intervening gap of 5 or 60 days limited to home cage. Correlations without ≥ 5-day break in training (light). (D) Model for computing expected field number correlations in (A) assuming each cell’s (relative) propensity was perfectly preserved across sessions. Propensities were drawn from gamma distributions (fit to each session) and modeled as order-preserving (i.e. each cell sits in same place in the CDFs across sessions). Each cell’s number of fields was modeled as a Poisson distribution based on its propensity in each session, yielding a joint distribution of numbers of fields per cell across sessions, from which expected correlation was computed. (E) Difference between observed field number correlation and expected correlation assuming perfect order preservation of propensity. Each group’s correlations were compared to 0 (perfect preservation). *p < 0.05, **p < 0.01, ***p < 0.001.

### Propensity is preserved across long time periods without task experience

Experiments in smaller environments have shown that hippocampal activity in one session makes activity in a subsequent session more likely, but not if the time gap is a day or more (Cai et al., 2016). Therefore, we tested whether the persistent place field number correlation in large environments required daily experience exploring virtual environments in the intervening period. We temporarily stopped training and kept animals in their home cage for 5 or 60 days without any virtual environment experience. Field propensity was highly correlated even across a 2-month gap, and the correlation across a given time interval was the same regardless of whether there had been a 5-day home cage-only period or not (Figure 4C).

### Time course of preservation of field propensity

We developed a statistical model to further investigate the time course of propensity. The place field number correlations decayed over time for both same and different environments (from ~0.6 for short time gaps to ~0.4 beyond a month) (Figure 4A). Since field occurrence is well-modeled as a spatial Poisson process (Rich et al., 2014), the same underlying field propensity would be expected to produce varying numbers of place fields in each session, at least for different environments. Thus, the correlation would be expected to be below 1 even if propensities were perfectly preserved. Additionally, the overall gamma propensity distribution exhibited some session-to-session variation (Figure 4D), which would also lower the correlation even if the ordering of each cell’s propensity with respect to the population were perfectly preserved. We created a model that took these factors into account by fitting a gamma distribution to each session, assuming the ordering of each cell’s propensity was perfectly preserved across sessions, then computing the expected field number correlation (Figure 4D). The difference between observed and expected correlations was used as a measure of how close the data was to a perfect preservation of (relative) propensities (Figure 4E). For different environments explored in the same session, correlations matched perfect preservation. While correlations gradually decreased below perfect preservation over a month, overall this analysis revealed each cell’s propensity was highly preserved across environmental contexts and time.

### Increased place field density at consistently rewarded locations

Salient stimuli such as reward are known to influence hippocampal representations, so we performed additional experiments to incorporate this important feature into our model. In particular, we manipulated reward contingencies to investigate the relationship between reward-related activity and place field propensity. In the experiments above and the previous large-environment study of propensity in freely moving animals (Rich et al., 2014), reward was delivered at random locations differing on each lap (Figure 5A). The overall density of CA1 place fields across the population is ~2-3x higher in consistently rewarded locations (Hollup et al., 2001; Dupret et al., 2010; Danielson et al., 2016; Zaremba et al., 2017; Gauthier and Tank, 2018; Sato et al., 2018), thus the propensity increases for at least some neurons. Is the increase concentrated in a small subset of cells that strongly and specifically respond to reward (Gauthier and Tank, 2018; Sato et al., 2018), or spread out across the population and, if so, in what manner (Figure 1C)?

**Figure 5.**
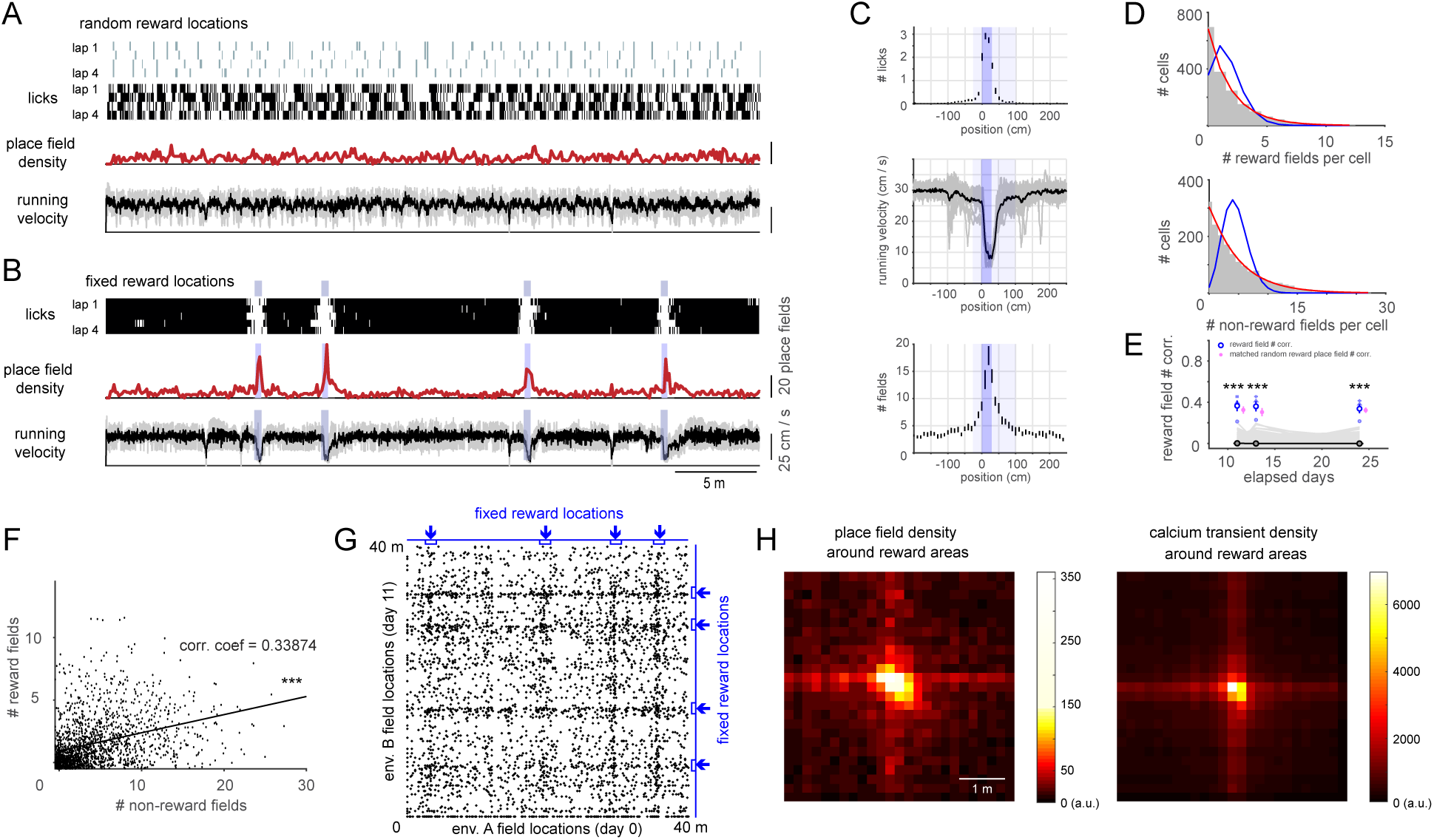
Hippocampal reward-related activity of each cell is correlated with its non-reward place field propensity. (A) Water reward and lick locations, place field density, and running speed (gray: each lap, black: average) in random reward sessions (Figures 1-4). (B) Same as (A) for sessions in which reward was provided at 4 fixed locations on every lap. (C) Licking, running speed, and field density around fixed reward locations (average of 4 reward locations per session, 3 sessions per animal). Dark blue: location where reward delivered; light blue: region of elevated field density defined as reward area used in subsequent analysis. (D) Reward field and non-reward field number per cell distributions. Equal Poisson (blue) and gamma-Poisson (red) fits. Field numbers combined across 3 fixed reward sessions, 4 mice. (E) Reward field number correlations (blue) across 3 distinct fixed reward environments (gray: shuffled data). Values were similar to place field number correlations (pink) for subsections of random reward environments matched for having the same mean number of fields (p = 0.49-0.83). (F) Scatterplot of number of reward and non-reward fields per cell (field numbers combined over 3 sessions, pooled over 4 mice). (G) Example scatterplot of field locations of individual cells in pair of fixed reward (blue arrows) environments (one point for every possible pair of session 1 and session 2 fields of each cell). See text for details. (H) Field or calcium transient pair density around fixed reward areas (±2.5 m around reward locations, 20 cm bins), summed over all pairs of fixed reward locations (e.g. the 16 intersection points in (G)) across all fixed reward session pairs. ***p < 0.001. See also Figure S4.

We repeatedly delivered reward at only 4 locations in every lap while preserving the total amount of reward per lap given in the random reward sessions. We presented 3 sets of 4 fixed reward locations (1 set in environment B, 2 in C) per animal (Figure S4A). For each set, we analyzed the last session after animals were exclusively trained in only one environment with one reward configuration for least 5 days in a row. Animals learned these high value locations – lick rate increased and running speed decreased in a predictive manner before each fixed reward location (Figures 5B and 5C). We observed the ~2-3x increase in field density around each rewarded location seen in the above studies (Figures 5B and 5C). The average field density was elevated 25 cm before to 75 cm after each 25 cm-wide rewarded region (Figure 5C), which we used as the reward areas (125 cm for each of 4 areas) in our analysis.

### Reward field propensity is highly skewed and preserved across distinct environmental contexts and time

We asked if, as with field propensity, cells differed in the probability of representing rewarded locations. To increase the statistics, we merged the number of reward fields (i.e. place fields in reward areas) per cell across the 3 distinct fixed reward sessions. The reward field number distribution was skewed, with most cells having few or no reward fields (Figure 5D). As in the random reward condition, the non-reward field (i.e. place fields outside of reward areas) number distribution was also highly skewed toward zero (Figure 5D). Both distributions were better fit by a gamma-Poisson than a Poisson distribution (Figure 5D).

We observed, as with field propensity, that each cell tended to display a similar number of reward fields across different environments and time (Figure 5E). Correlation values for non-reward fields did not differ from random reward condition correlations (p = 0.36-0.74, Figure S4B). While the reward field number correlation was lower than for non-reward fields, this could be explained by the higher variability associated with the smaller mean number of reward fields. Indeed, place field number correlations for subsections of random reward environments matched to have similar field numbers (25% of total length, mean place field number = 0.51 versus mean reward field number = 0.52) showed similar values to reward correlations (Figure 5E, pink, p = 0.49-0.83 for 3 matched time gaps).

### Reward field propensity is correlated with field propensity

Since both field propensity and reward field propensity were skewed and stable across environments and time, we investigated the cell-by-cell relationship between these propensities. Do cells with higher field propensity also tend to respond more at rewarded locations? Or are these properties uncorrelated? Or is a there a subset of cells (~5%) specialized for representing reward (Gauthier and Tank, 2018; Sato et al., 2018)? For this analysis, we combined reward and, separately, non-reward field numbers of each cell across the 3 distinct fixed reward sessions and pooled across animals.

First, reward and non-reward field numbers of individual neurons were correlated with each other (Figure 5F) (r = 0.34, p = 7.7×10^-48^, 1735 cells tracked across all 3 fixed reward sessions; mouse 1-4: 522, 451, 616, 146 cells).

Next, we looked for evidence of a class of reward-specific neurons. Most cells (90.3%) that had a reward field also had one or more non-reward fields. No cell had 12 reward fields (a field at every reward location in the 3 sessions) and 0 non-reward fields; only 1 cell had 12 reward fields, and it also had 8 non-reward fields (Figure S4C). The 5% of cells with the most reward fields exhibited many non-reward fields (Figure S4D). 5.8% of cells had at least 1 reward field but 0 non-reward fields. The reward field number distribution of these cells was highly skewed towards 1 reward field out of 12 possible (Figure S4E). Only 1 cell exhibited context-specific reward-specific activity (i.e. either 4 or 0 fields in the 4 rewarded locations in each of the 3 fixed reward configurations, Figure S4F). These results are consistent with reward activity representing a conjunction of reward × place, rather than reward per se.

We also investigated which cells contributed to the overall increased field density at the rewarded locations. For each pair of fixed reward sessions, we created a 2D scatterplot that contains a point at (x,y) whenever a neuron was observed to have a (reward or non-reward) field at location x in session 1 and location y in session 2 (Figure 5G) (Gauthier and Tank, 2018; Sato et al., 2018). Cells with n_1_ fields in session 1 and n_2_ fields in session 2 thus contributed n_1_ × n_2_ points. (If a cell had 0 fields in a session, point(s) were placed at 0 for that session.) The increased density of points along vertical and horizontal lines corresponding to reward locations in a session indicated that cells contributing to the excess reward field density (over the density in non-rewarded areas) also had non-reward fields (Figure 5G). This was also visible in the average of the scatterplots centered on every point corresponding to a rewarded location in both sessions (Figure 5H). That is, excess reward field density did not exclusively emerge from cells that only had reward fields. The result was the same when considering all calcium transients irrespective of whether they occurred in a field (Figure 5H).

### Reward multiplies each cell’s field propensity by a constant gain factor

Since we discovered a clear correlation between a cell’s reward and non-reward field propensities, we thus examined several statistical models of the joint distribution of reward and non-reward fields of individual cells across the population. The simplest model that would give correlated reward and non-reward field numbers is a “constant gain” model, in which reward field propensity equals a common gain value times the non-reward field propensity. The observed joint distribution of reward and non-reward field numbers was well-accounted for by the perfectly correlated (i.e. perfect order preserving) constant gain model in 2 of the 4 animals (Figure 6A, right bottom 3 panels). We expanded the constant gain model by using a copula to generate correlated field propensities in reward and non-reward regions (Figure 6A). The use of a copula guarantees the separate (marginal) reward and non-reward field number distributions are preserved while allowing variation (in this case with a single continuous parameter) between perfectly correlated and completely uncorrelated reward and non-reward field propensities. The copula parameter was fit to the reward and non-reward field number correlation (i.e. not to the full joint distribution). Goodness of fit of the resulting model was assessed by estimating the similarity (Jensen-Shannon divergence, D_JS_) between the model and full joint distribution of reward and non-reward field numbers. For all animals, these copula models fit the data with high correlations (mean = 0.75) between reward and non-reward propensities (Figure 6B). We tested alternative models in which the gain could vary nonlinearly but monotonically with the underlying (non-reward) propensity. The copula models were better (i.e. lower D_JS_ from the data) than these more complex models that notably had more parameters which were estimated from the full joint distribution itself (Figure 6C). Thus, the representation of value in the hippocampus can be largely explained by a uniform value-driven scaling of each cell’s underlying place field propensity.

**Figure 6.**
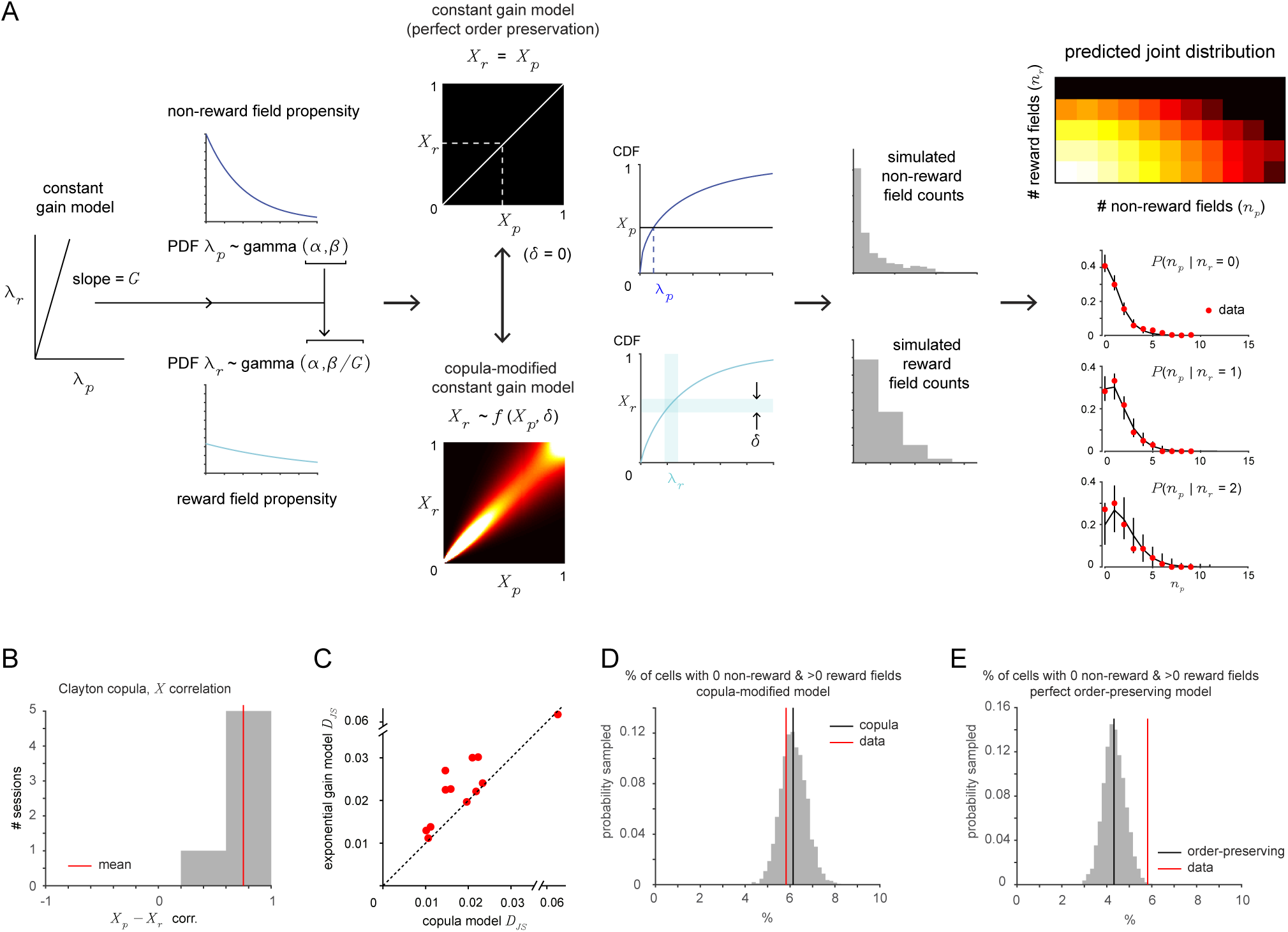
Reward-related activity is proportional to non-reward field propensity. (A) To model reward activity, we assumed the distribution of reward propensities across cells had a constant gain with respect to the non-reward field propensity, yielding a scaled gamma distribution. We tested models with perfect order preservation (each cell has reward propensity of a constant times its non-reward field propensity) and parameterized correlated activity (copula), where reward and non-reward field numbers are determined by Poisson processes with rates given by the respective propensities. This yields joint distributions of reward and non-reward field numbers per cell that can be compared to data. (B) Correlations of copula models fit to each fixed reward session were very high for most sessions (value of 1 implies perfect order preservation). (C) Comparison of performance of copula model and a parameterized gain model in predicting actual joint distribution for each fixed reward session, showing consistently better agreement (lower Jensen-Shannon divergence) of copula model despite fitting 1 fewer parameter (4 instead of 5). See text. (D) Distribution of proportion of “reward only” cells predicted by copula model, showing the observed number of reward-only cells is well-predicted by the copula model. (E) Same as (D), but for order-preserving model, showing most of the observed cells are accounted for by the perfect order-preserving model.

Using the constant gain model, we revisited the question of whether there is a subset of reward-specific cells. We computed the predicted distribution of cells with at least 1 reward field and 0 non-reward fields, i.e. candidate reward-specific cells, based on the model. The copula model could account for all (Figure 6D) of the 5.8% of cells observed to have at least 1 reward field and 0 non-reward fields, and the perfect order preserving model could account for most of them (median = 4.3%, 95% CI = 3.4-5.3%, Figure 6E). Both models, by definition, assume no reward-specific subpopulation.

### Propensity is anatomically distributed in a salt-and-pepper fashion

Since propensity was a preserved property of individual cells under all conditions tested, we checked whether there was any anatomical organization of propensity (Figure 7A). Adjacent cells had widely different numbers of fields (Figures 7B and 7C). There was a small correlation in the field numbers of nearby cells (all pairs of cells with centers <25 µm apart; mouse 1: r = 0.18, p < 0.001, 737 pairs; mouse 2: r = 0.035, p = 0.49, 380 pairs; mouse 3: r = 0.089, p = 0.015, 753 pairs; mouse 4: r = 0.14, p = 0.061, 171 pairs). However, when we controlled for the moderate gradient in propensity across the superficial-deep (Mizuseki et al., 2011; Danielson et al., 2016) and proximal-distal axes (Henriksen et al., 2010) described above by using a low-pass spatial filter (i.e. subtracting from each cell’s field number the mean field number within an 80 µm radius), the correlation in field numbers of nearby cells disappeared (Figure 7D). Therefore, propensity is organized in a largely salt-and-pepper manner.

**Figure 7.**
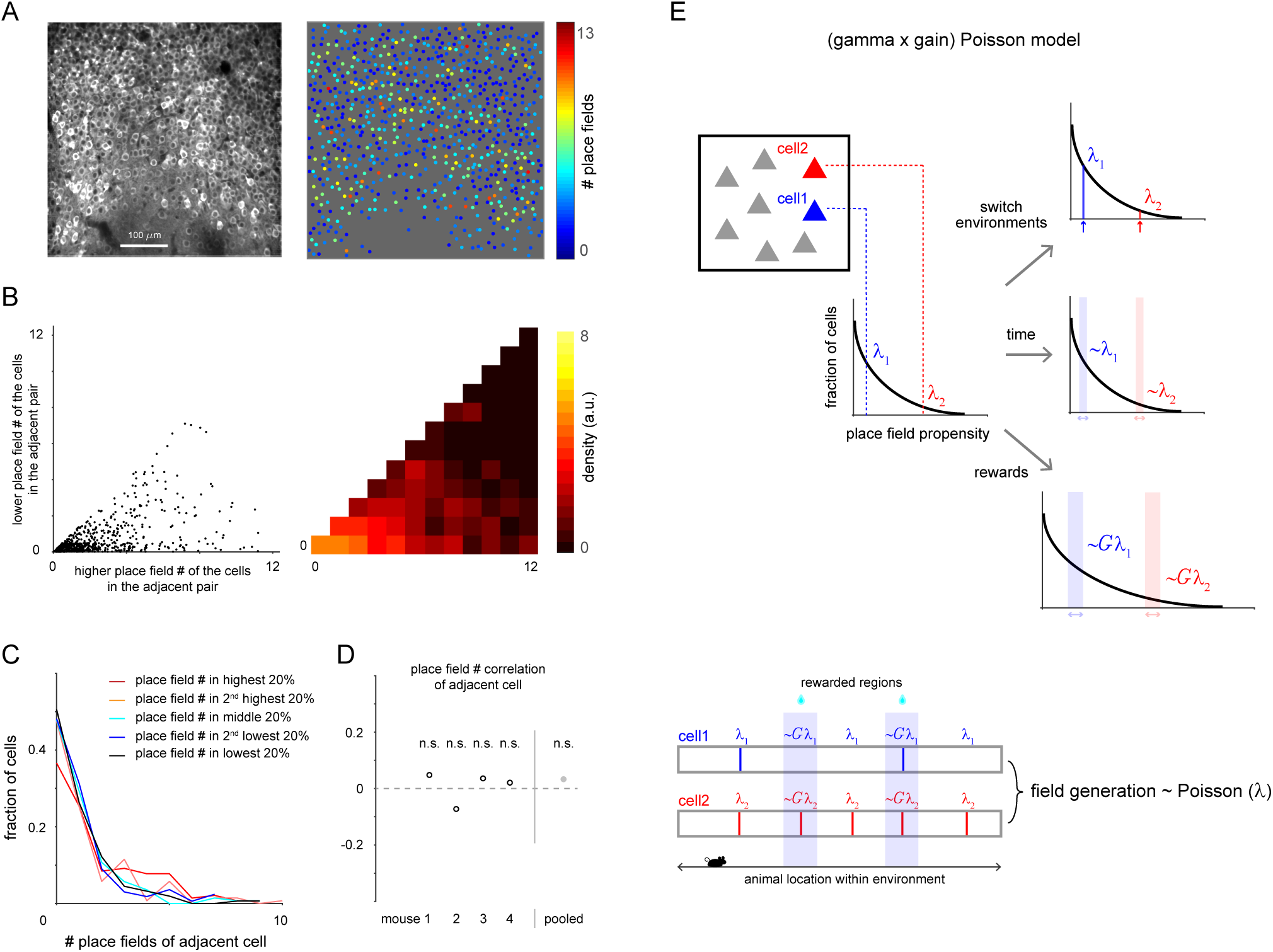
Cells with widely different propensities are anatomically arranged in a salt-and-pepper fashion. (A) Standard deviation image of GCaMP6f fluorescence signal and place field numbers of corresponding neurons. (B) Scatterplot (jittered, left) of number of fields of each pair of cells with centers <25 µm apart, and corresponding density (right) (mouse 1), showing adjacent cells have widely different field numbers. (C) Distribution of number of fields of cells <25 µm away from cells with field numbers in a given propensity quintile (mouse 1). (D) Correlation of number of fields between pairs of cells <25 µm apart for each mouse and pooled (right). Number of fields of each cell was corrected to control for known propensity gradient across the anatomical axes. See text. (E) Schematic of “(gamma × gain) Poisson” model of hippocampal representations based on the findings in this study.

### Unified model of hippocampal representations

Bringing together the above results, we can describe at a statistical level how the hippocampus will represent spatial contexts under a wide range of conditions with a simple generative model we call the “(gamma × gain) Poisson” model (Figure 7E). Specifically, for each cell, one assigns it a propensity randomly selected from the gamma distribution of propensities, corresponding to its salt-and-pepper distribution. Given its propensity, each cell will randomly express place fields in a spatially Poisson manner, with the propensity being the Poisson rate parameter (Rich et al., 2014). At rewarded locations, every cell’s propensity will be multiplied by a common gain factor. Finally, each cell’s propensity predicts its activity independent of environmental context and is highly preserved for a month or more. The key parameter underlying all aspects of the model is the cell’s propensity. Modifications to this model to account for additional details would include a small shift of the gamma distribution to slightly lower (higher) mean propensities for more superficial (deep) and more proximal (distal) regions of CA1 (Henriksen et al., 2010; Mizuseki et al., 2011; Danielson et al., 2016), a small deviation of reward propensity from a constant gain times the non-reward propensity, and a small drift of propensity values over time.

## Discussion

Here we have shown that a single function – the field propensity distribution – with a simple shape (a 2-parameter gamma distribution (Rich et al., 2014)) underlies many fundamental features of hippocampal activity. The propensity of each cell determines how many place fields that cell will have and how likely it will respond at locations consistently associated with reward, applies to any environment, and is a stable property of the cell over time. In the absence of location-specific rewards or other salient stimuli, a cell’s fields are randomly located (i.e. spatial Poisson) given its propensity (Rich et al., 2014), thus propensity is the key parameter determining the structure of hippocampal representations. Furthermore, we show that propensity is anatomically distributed in a largely salt-and-pepper manner, which therefore imposes a physically dispersed organization onto this representational structure. All of these features can be captured by a compact model, the “(gamma × gain) Poisson” model (Figure 7E). In particular, the results reveal persistent, order-of-magnitude differences (Figures S3B-S3D) between neighboring CA1 pyramidal cells in the feature these cells are most well-known for, i.e. having place fields (including fields in rewarded locations). That is, the probability that a given neuron will participate in the representation of any given location, context, or other item (e.g. a rewarded location) greatly and consistently varies across cells, with some cells being active in the representations of many items and others in only a very select subset of them. These large, persistent differences strongly shape the hippocampal representation of any given environmental context and thus very likely impact how the spatial code – and presumably the code for episodic memory in the human hippocampus (Gelbard-Sagiv et al., 2008) – is encoded, read out, or associated with other behaviorally relevant information.

Previous work had shown that (1) the average firing rates of hippocampal cells are skewed and correlated across sleep/wake states and across environments within a day (Hirase et al., 2001; Mizuseki and Buzsaki, 2013), but that (2) activity levels change over a timescale of a few hours to a week (Ziv et al., 2013; Rubin et al., 2015; Cai et al., 2016). Here, tracking the same cells over a month or more in multiple large environments enabled us to determine that propensity is highly preserved across environments and time. Imaging also allowed us to determine the high-resolution anatomical organization of propensity. Finally, unlike previous work, we studied propensity and reward activity together, and discovered they are closely related.

The preservation of each cell’s propensity across environmental contexts with distinct visual configurations as well as (its relative propensity) in rewarded locations and the dark (Figures 3H and 3I) suggests a modified view of the origin of receptive fields such as place fields. Receptive field models have generally focused on the number and strength of individual inputs as determining the output of a cell. Such a view can be expressed as “Output ~ Input”, with variations in excitability being small relative to variations in inputs. The critical role of inputs has been demonstrated in experiments showing that sensory cues can control place field location (O’Keefe and Conway., 1978; Muller and Kubie., 1987; Deshmukh and Knierim, 2013; Geiller et al., 2017). However, while our findings are consistent with the role of inputs in determining tuning (i.e. field locations), they suggest that whether tuned inputs ultimately lead to output (i.e. field number) strongly depends on a global, excitability-based property of each postsynaptic cell. That is, “Output ~ Input × Excitability”, with large variations in excitability controlling the level of output.

This picture is supported by intracellular recordings of place and silent cells in behaving animals. These recordings showed that hippocampal neurons differ in intrinsic excitability (Epsztein et al., 2011), that place field activity is correlated with excitability (Epsztein et al., 2011; Cohen et al., 2017), and that manipulating excitability controls whether spatially tuned inputs are converted into place field spiking (Lee et al., 2012; Hsu et al., 2018). These recordings also revealed that inputs into hippocampal neurons are spatially tuned across the full extent of the environment, with multiple peaks of the subthreshold membrane potential as a function of location independent of whether the cell is active or not (Cohen et al., 2017; Hsu et al., 2018). Such tuning (Lee et al., 2012; Cohen et al., 2017; Hsu et al., 2018) therefore provides each cell with a large number of potential place fields from which any number can be converted into output depending on the neuron’s excitability. The spatial Poisson-like randomness of field locations would arise from effectively thresholding the sum of the large number of spatially tuned inputs to the cell. Intracellular and extracellular recordings also showed that the number of place fields a cell has in a novel environment is correlated to its firing pattern or rate during a preceding period of anesthesia or sleep, before any exposure to inputs from the environment, again consistent with excitability controlling propensity (Epsztein et al., 2011; Rich et al., 2014). Consistent with this, extracellular recordings have shown correlated firing rates across successive sleep and wake states (Hirase et al., 2001; Mizuseki and Buzsaki, 2013). Alternatively, differences in the total number, strength, and/or distribution (Druckmann et al., 2014) of inputs to each cell could contribute to differences in propensity. However, place and silent cells had different bursting patterns in response to intracellular current injection during a period when inputs were inactive (immediately after obtaining the whole-cell recording, before residual low-calcium internal solution was cleared from the cell’s neighborhood) (Epsztein et al., 2011), supporting a cell-intrinsic basis of propensity. In addition, multiple approaches have demonstrated the role of excitability in determining which subset of cells will participate in fear memory representations in the amygdala (Han et al., 2007; Silva et al., 2009; Zhou et al., 2009; Rashid et al., 2016) and hippocampus (Cai et al., 2016).

What mechanisms could underlie the order-of-magnitude range and gamma distribution of propensities across neighboring neurons that have generally been considered to constitute a single cell type (i.e. dorsal CA1 pyramidal cells at the same superficial-deep and proximal-distal location)? Electrophysiological studies have shown differences in excitability and other cellular or local circuit properties of hippocampal pyramidal neurons (Epsztein et al., 2011), especially along the proximal-distal and superficial-deep axes (Jarsky et al., 2008; Graves et al., 2012; Lee et al., 2014, Valero et al., 2015) axes. The ~30-40% differences in propensity reported along the proximal-distal (Henriksen et al., 2010) and superficial-deep (Mizuseki et al., 2011; Danielson et al., 2016) axes could potentially be explained by these gradients. However, the wider range and salt-and-pepper distribution of propensities observed here imply larger fluctuations in properties between adjacent cells than suggested by such gradients. The persistence and apparent cell-intrinsic basis of propensity differences suggest transcriptomic differences between CA1 pyramidal cells, which in turn could arise from developmental differences (Zeisel et al., 2015; Cembrowski et al., 2016, Mallory and Giocomo, 2018; Soltesz and Losonczy, 2018). In particular, differences in specific conductances (Hsu et al., 2018) could underlie excitability-based propensity differences. Morphological differences may also contribute (Diamantiki et al., 2016). Reward-based increases in propensity could arise from amplification by entorhinal cortex inputs (Jarsky et al., 2005; Dudman et al., 2007; Bittner et al., 2015; Basu et al., 2016) of CA3 inputs, as CA3 does not show increased field density in rewarded locations (Dupret et al., 2010).

What could be the function of the persistent, gamma distribution of propensities shown here? As mentioned above, the significant fraction of cells with consistently low propensity would result in a sparse code for location, including for rewarded places, which could be used for goal-directed navigation. However, while sparse codes have known value (Tsodyks and Feigelman, 1988; Olshausen and Field, 2004; Katkov et al., 2017), what would be the value of the “dense code” among the subset of cells with consistently high propensity? One possible function of having a wide range of propensities may be to ensure that enough cells can participate in distinguishing environments of any size (Rich et al., 2014). For instance, for a small environment, high propensity cells would ensure that the engram (Liu et al., 2012) consisting of all cells with fields in the environment would not be proportionately small. Recent work has shown that CA1 cells that learn novel environment structure tend to have fewer fields (Grosmark et al., 2016). More generally, the persistent, continuous distribution of propensities allows for the simultaneous existence of a sparse code, dense code, and everything in between among pre-specified subpopulations of neurons.

We could explain the increase in field density around rewarded locations as resulting simply from multiplication of each cell’s field propensity by a constant gain. Other work has instead attributed the increase to a small subset (~5%) of cells that are active only at rewarded, and not at non-rewarded, locations (Gauthier and Tank, 2018; Sato et al., 2018). Our data did contain a fraction of cells (~5%) that had at least one reward field and no non-reward fields, but our model could account for them as cells that could have had non-reward fields but did not by chance (though we cannot rule out the possibility of a very small subset of reward-specific cells). Also, in contrast to the other work (Gauthier and Tank, 2018), the excess field density we observed at rewarded locations could not be accounted for by cells only activated by reward (Figures 5G and 5H). Thus, while previous work argued that reward responses, and therefore reward input, is specific to a small subset of cells, our results are consistent with all cells receiving reward input. The different results could be due to our use of 3 large, ~40 m environments with 4 rewarded locations each, compared to the use of 2 smaller (~2-4 m) environments each with a single rewarded location in the previous studies.

We could explain the variability in the number of fields per unit track length each cell had across environments and time as resulting from a fixed propensity plus the expected Poisson noise, as well as some global modulation of the entire propensity distribution, and a small amount of drift over time. Previous work employing smaller (~1 m scale) environments has shown evidence that propensity changes over a timescale of hours to a week using a binary measure of propensity, i.e. whether the cell was a place or silent cell (Ziv et al., 2013; Rubin et al., 2015; Cai et al., 2016). The difference could potentially be explained by our use of larger environments, first because of differences in the animal’s experience and encoding, and second because the larger range of field numbers allow for a higher resolution measure of propensity than binary assessment. The previous work also showed an elevated preservation of propensity between pairs of experiences occurring within a few hours (Cai et al., 2016), attributed to activity-dependent mechanisms that transiently increase excitability in recently active cells (Han et al., 2007; Silva et al., 2009; Zhou et al., 2009; Cai et al., 2016; Rashid et al., 2016). Consistent with this, our data showed a higher preservation of propensity across environments explored on the same day (Figures 4A and 4E). The dominant trend we found, though, was preservation of propensity with minor degradation over a month (Figure 4E), and preservation across periods of days to months in a home cage without task experience (Figure 4C).

The large and persistent differences in propensity we observed among nominally similar cells could be a general organizing principle for other brain areas, especially those that learn distributed representations of arbitrary internal or external stimuli (Rigotti et al., 2013). In addition to CA1 (Rich et al., 2014), large differences in propensity have been reported in other hippocampal areas (Alme et al., 2014; Witharana et al., 2016). A highly skewed distribution of propensities has also been observed in the fly mushroom body, where many Kenyon cells responded to only a few odors and a few to many odors (Turner et al., 2008). Given a particular fixed distribution of propensities across the population, memory capacity (in terms of the total number of distinct representations) would be maximized if each cell was randomly selective for stimuli at a rate constrained by its propensity. This has been shown for CA1 pyramidal cells, which have place fields in random locations (i.e. spatial Poisson with propensity as the rate parameter) (Rich et al., 2014). Generalizing to any brain area, an advantage of having many persistently low propensity neurons would be the resulting sparse code for whichever stimulus space is represented. Having a subset of persistently high propensity neurons could, as mentioned above, ensure enough cells represent each stimulus, and could facilitate associating multiple representations via overlap.

## Author contributions

J.S.L. and A.K.L. designed the experiments. J.S.L. performed the experiments. J.S.L. and J.B. performed the data analysis with input from A.K.L. and S.R. J.B. and S.R. constructed the models with input from A.K.L. and J.S.L. A.K.L., J.S.L., J.B., and S.R. wrote the paper.

## Acknowledgments

We would like to thank Nick Sofroniew, Karel Svoboda, Vasily Goncharov, Igor Negrashov, Spencer Taylor, Jon Arnold, Jason Osborne, Bill Biddle, and Bruce Bowers for technical assistance with imaging and constructing the experimental apparatus and components, and Mark Bolstad and Jeremy Cohen for technical assistance with the virtual reality software and hardware. This work was supported by the Howard Hughes Medical Institute.

---

**Figure S1.**
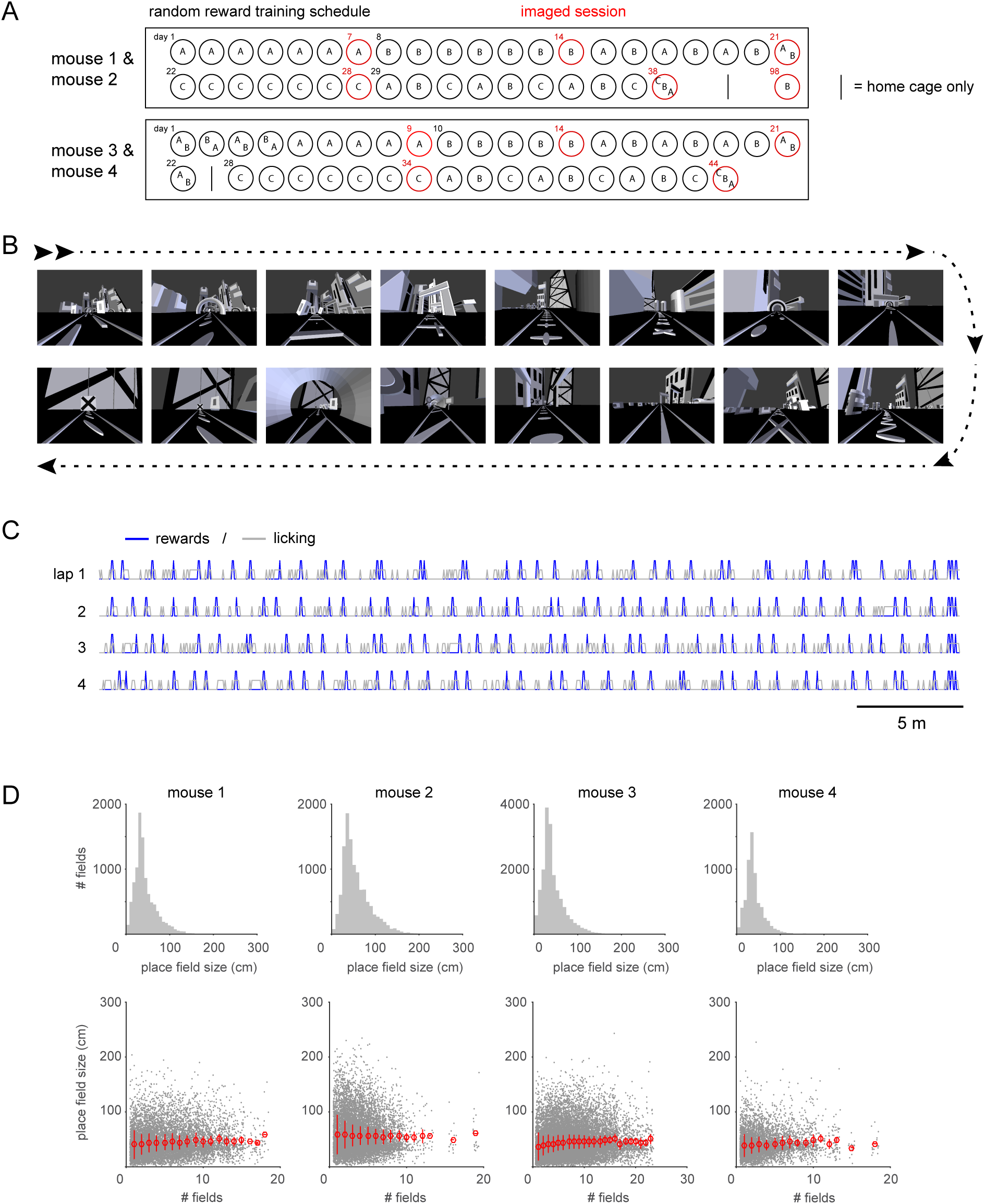
Training schedule, example environment, behavior, and place field size in virtual environments, Related to Figure 1. (A) Daily training schedule for mouse 1 and mouse 2 (top), and mouse 3 and mouse 4 (bottom). A, B, C represent the 3 environments explored by each animal (selected from 4 environments), where A represents the first environment explored by each animal regardless of its identity. Red circles indicate the calcium imaging sessions. (B) Sequential view along the running path for one of the 4 virtual environments. (C) Mouse licking behavior in each lap. Blue lines indicate randomly delivered water rewards and gray lines indicate mouse licking. (D) Distribution of place field size (top). Place field size (44 ± 25 cm, 57 ± 32 cm, 44 ± 26 cm, 40 ± 21 cm in each mouse, mean ± s.d.) was not correlated with the number of place fields of the corresponding cell (bottom). Average place field size for cells with a given number of place fields (red, mean ± s.e.m.).

**Figure S2.**
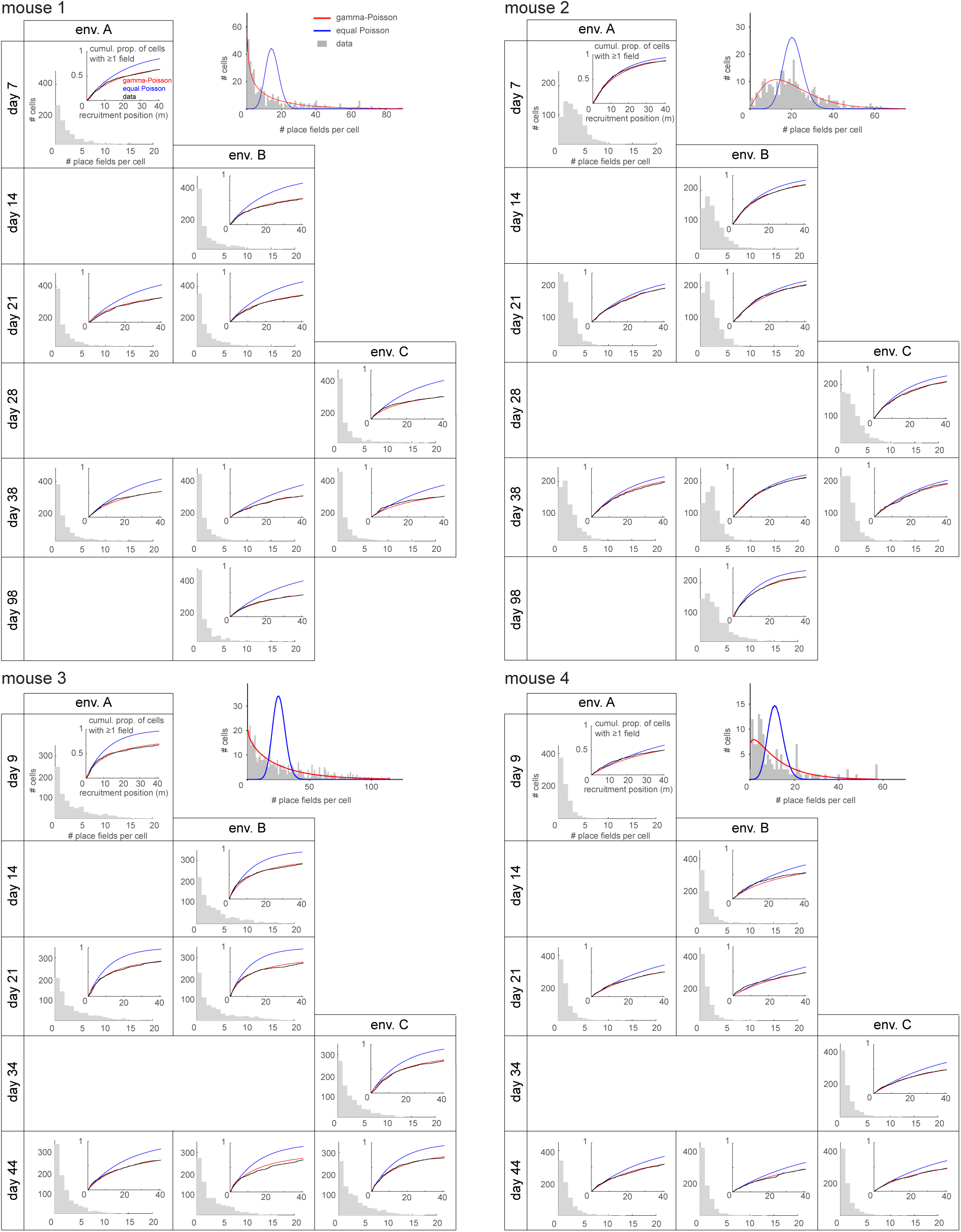
The number of place fields of each cell in different virtual environments on different days is well-described by the gamma-Poisson model, Related to Figure 1. Each panel shows the distribution of the number of place fields of each cell in each (random reward) session for the 4 mice. Each mouse had somewhat different distributions, but across the sessions of a given mouse the distribution was preserved over time and environments (distributions on days 7, 14, and 28 for mouse 1 and 2, and 9, 14, and 34 for mouse 3 and 4 were used for these comparisons, K-S test, p > 0.15). Inset in each panel shows the cumulative recruitment of cells along the track (i.e. a cell is defined as recruited by position x if it has a place field between 0 and that position) (black) with the prediction from the equal Poisson model (blue) and the gamma-Poisson model (red). This shows good agreement with the gamma-Poisson model. For each cell with ≥6 fields in a session, the distribution of field locations and distribution of inter-field intervals did not differ from the spatial Poisson model predictions of uniform and exponential, respectively (Anderson-Darling test, 0/4264 cells with p < 0.05, adjusted for false discovery rate, q = 0.25), consistent with the fields of each cell being distributed randomly in space, as shown previously (Rich et al., 2014). Top right plot for each animal shows the distribution of the total number of place fields across all (random reward) sessions per cell (sum over the 8 or 9 sessions shown in the bottom left panels).

**Figure S3.**
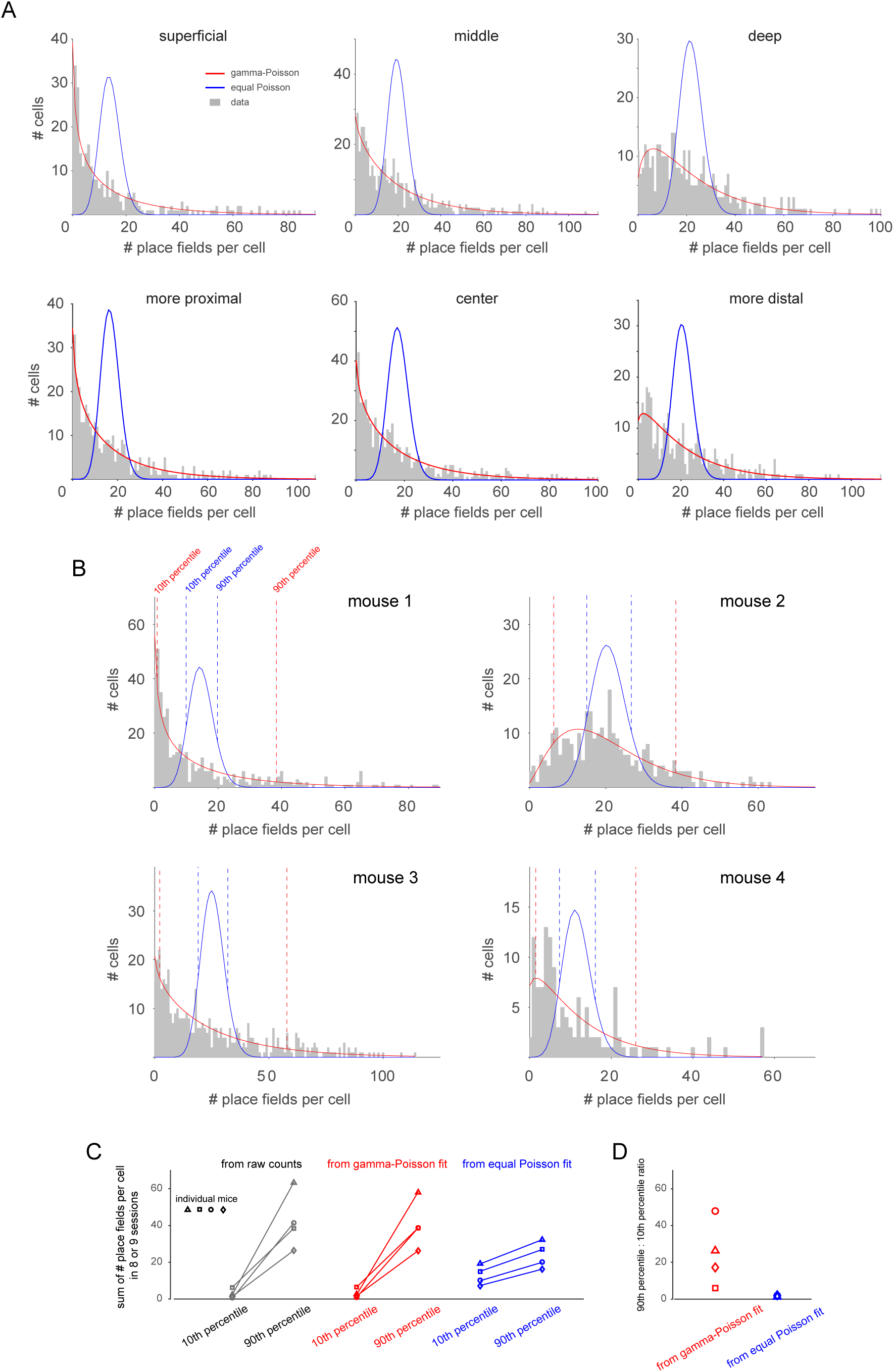
The distribution of the number of place fields per cell varies by an order of magnitude across the population, Related to Figure 1. (A) The distribution of the total number of place fields across all (random reward) sessions per cell for subpopulations of cells along the superficial-deep (top) or proximal-distal axes (bottom) (pooled data from the 4 mice). Superficial-deep location of each cell was estimated from a z-stack of images taken above and below the imaging plane. The cells from 0-35% of the way from the superficial to deep edges of the pyramidal cell layer were included in the superficial group, and from 65-100% in the deep group. The cells from 40-60% were included in the middle group. The different sizes of the ranges were used to balance the number of cells in each group. For the proximal-distal axis, we divided our imaging plane into 3 regions: 1/3 of anterior lateral region as proximal, 1/3 of posterior medial region as distal, and the rest 1/3 as center (our imaging plane is the left hemisphere dorsal CA1). Equal Poisson (blue) and gamma-Poisson (red) probability density functions were fit in each panel. The large deviation from the equal Poisson fit demonstrates that cells do not have the same field propensity. (B) The distribution of the total number of place fields across all (random reward) sessions per cell (gray), with equal (blue) and gamma-Poisson (red) fit (same as in Figure S2). Dotted lines indicate 10th and 90th percentile of each fit of corresponding color. (C) Comparison of 10th and 90th percentile of total number of place fields across all (random reward) sessions for each animal (unique symbol per mouse). (D) The ratio of 90th:10th percentile propensity from gamma-Poisson (red) and equal Poisson (blue) fit.

**Figure S4.**
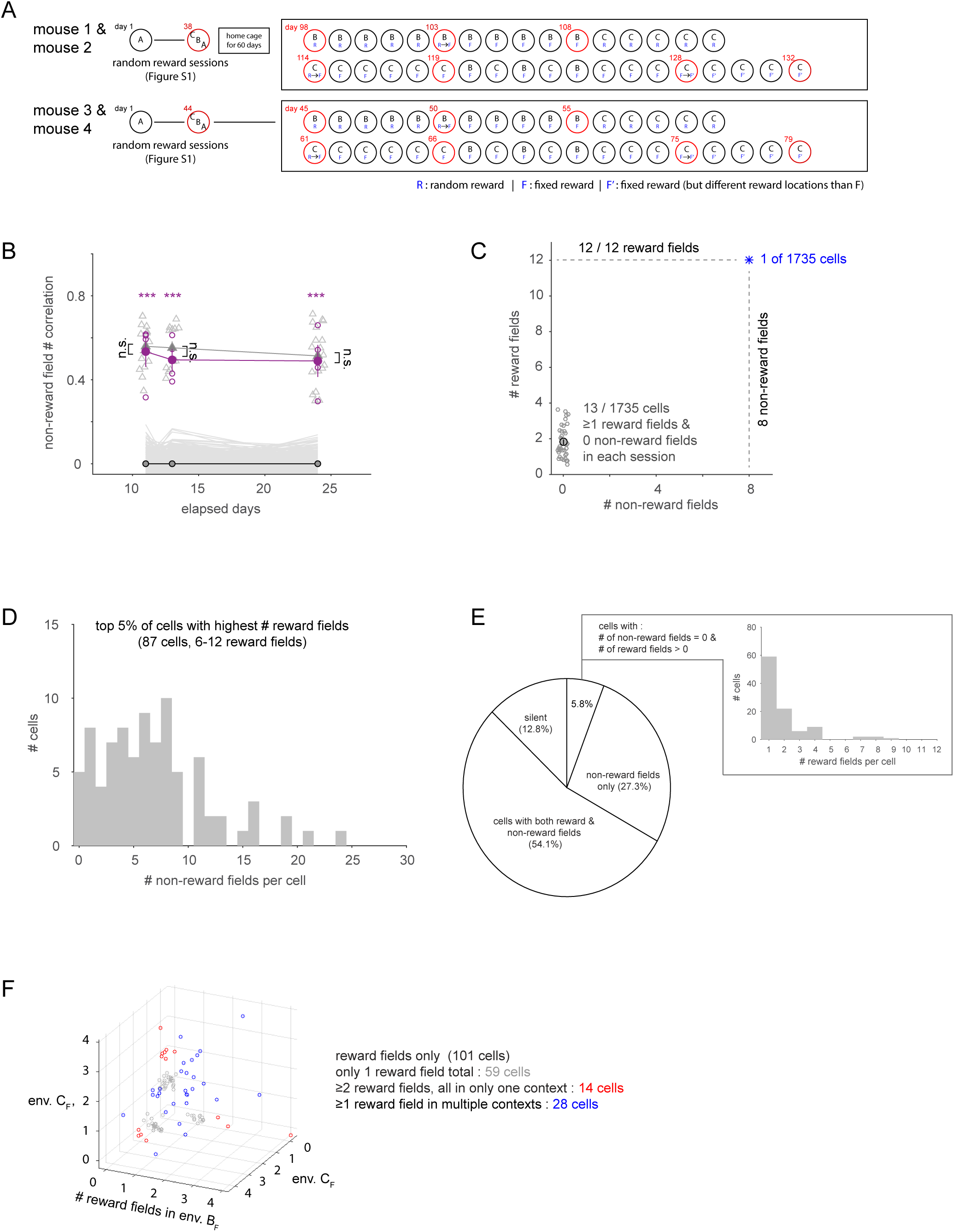
Reward and non-reward activity in fixed reward location experiments, Related to Figure 5. (A) Daily training schedule for mouse 1 and mouse 2 (top), and mouse 3 and mouse 4 (bottom). Red circles indicate the calcium imaging sessions. C_F_ and C_F_’ correspond to the same environment but with a different set of 4 fixed reward locations. (B) The number of place fields per cell in non-rewarded regions of the environments in the fixed reward sessions were correlated across environments and time (magenta filled: mean ± s.e.m., magenta open: individual session pairs; ***p < 0.001 versus shuffled data: gray lines and black circles for means at bottom) and not different from the correlations of field numbers in the random reward sessions (gray triangles filled: mean ± s.e.m., open: individual session pairs). (C-F) Cells with reward-related activity have a spatial component as opposed to being only responsive to reward. Cell data was pooled across the 3 fixed reward sessions of the 4 mice (BF: day 108 for mouse 1 and 2, day 55 for mouse 3 and 4, C_F_: day 119 for mouse 1 and 2, day 66 for mouse 3 and 4, C_F_’: day 132 for mouse 1 and 2, day 79 for mouse 3 and 4), using only those cells tracked across all 3 sessions: 1735 cells total. (C) 13 of 1735 cells had at least 1 reward field in every fixed reward session and no non-reward fields. In each environment (13 × 3 sessions, bottom left), these cells generally had reward fields in only a subset of the 4 rewarded locations. Only 1 cell had reward fields in all 12 rewarded locations across all 3 sessions, and it also had 8 non-reward fields. (D) The ~5% of cells with the largest total number of reward fields generally each had several non-reward fields. (E) The proportion of the 1735 cells in each of 4 categories (cells with both reward and non-reward fields, non-reward fields only, reward fields only, and those with no fields). 5.8% of cells (101/1735) had only reward fields, and these cells generally had fields in only a few of the 12 rewarded locations (box). 90.3% of cells (i.e. 54.1% / (54.1% + 5.8%)) that had a reward field also had ≥1 non-reward field. (F) To search for context-specific reward cells (i.e. cells with fields in every rewarded location in ≥1 environmental context and 0 reward fields in the other contexts), we plotted the number of reward fields in each of the 3 environments for the 101 cells (5.8% of cells) with reward fields only. These cells generally had fields in only a subset of the 4 rewarded locations in any given environment.

## Materials and Methods

### Transgenic mice and chronic window surgery

All procedures were performed in accordance with the Janelia Research Campus Institutional Animal Care and Use Committee guidelines on animal welfare. Four male GCaMP6f mice (Thy1-GCaMP6-WPRE transgenic mice (Dana et al. 2014), 8-10 weeks old) were used in all experiments in this study. Mice were anesthetized with isoflurane (1.5-2.0%) and a craniotomy was centered at −2.0 mm AP and 1.6 mm ML from bregma over the left hemisphere using a 3 mm diameter trephine drill bit (Trephine Bur, 3.0/2.0mm, THB30, Osung USA). The cortex and corpus callosum, but not alveus, above the dorsal hippocampus was removed by gentle suction with saline using a 27 gauge blunt-tip needle. After cortex / callosum removal, a glass coverslip (T 0.1 mm, D 3 mm, round, No.0, Warner Instruments, CS-3R-0) that was previously attached (using (Norland Optical adhesive 81) to the bottom of stainless steel cannula (OD 3.02 mm, ID 2.54 mm, L 1.75 mm, New England Small Tube Corp. Type 304, stainless steel, 11TW) was implanted over the dorsal CA1 region. A thin layer of Kwik-Sil (WPI) was applied between the cannula and skull. The top part of the cannula and custom headplate (stainless or titanium) were fixed to the skull using dental cement (Meta-bond) and dental acrylic. Body temperature was maintained at 37°C using a temperature controlled heating pad.

### Virtual reality and behavioral setup

We used our custom-made virtual reality (VR) software ‘mouseoVerR’ and spherical treadmill system which were described in detail previously (Cohen et al., 2017). Instead of using three projectors that projected onto the back of a screen with 225° field of view, in this study we used a single projector that projected off a flat then convex mirror onto the front surface of a 180° cylindrical screen. The vertical field of view was from 20° below to 60° above the horizontal plane through the animal. A horizontal 145°-wide rendered view image in mouseoVeR was projected onto the 180° screen and the image distortion from the convex mirror was compensated using a function in mouseoVeR. We used the blue channel of the projector only and, in addition, a blue filter was located in front of projector (Valley design, FG-B03_3). Licks were detected from beam breaks by the mouse’s tongue.

### Behavioral training

Four large virtual environments were made in Blender software. Each environment had a narrow ~14-cm-wide ~41 m-long winding path of different shapes that the mice were constrained to follow. Features of various spatial scales were placed around the environment, with smaller objects placed along the track to provide optic flow when running. The animal’s virtual view was constrained to be parallel to the track but they were allowed to move side-to-side within the width of the path. To help mice discriminate between the different environmental contexts, we also had a different, constant background odor cue (banana, cinnamon, lemon or apple) present for each environment. A unique combination of environment and odor cue was used for each mouse, but for a given mouse the combination was consistent throughout training. The mice underwent initial VR training using four different smaller environments (oval, circular, linear, linear with 15° turns, total track length ~185-325 cm) for habituation to head fixation and running on the treadmill through virtual environments. After 3 days of this smaller environment training, mice were trained in the large environments and in each session ran a total of 8-12 laps in 1-3 environments (i.e. 4-8 laps per environment). At the end of each lap, the animal was teleported back to the beginning of the environment’s track, the virtual view was locked at the start location for 5 s, then the animal was allowed to continue with the next lap. To make mice run consistently, mice were placed on water-restriction (1.2-1.4 ml/day) at least 3 days before starting initial VR training. For the random reward condition, 2 µL drops of water was delivered at 40 pseudorandom locations in each lap. For the fixed reward condition, we used the same environments except there were 4 fixed reward locations (and in each reward location 5 4-µL drops of water were delivered within a 25 cm segment of track). Supplemental water given based on the amount of water delivered in the VR training.

### Two-photon calcium imaging

Calcium imaging was performed with a custom-built two-photon microscope with a resonant scanner (Janelia Research Campus, MIMMS). The light source was a Chameleon Vision II (Coherent) laser tuned to 940 nm. The objective was a 16x water immersion lens (Nikon, 0.8 NA). Functional images were acquired using ScanImage (Vidrio) software. Imaging was done in one plane, the size of the imaging plane was 500 µm x 500 µm (512 x 512 pixels), and the sampling rate was 30 Hz. The same imaging plane was found on successive sessions by visual comparison with a saved image from the first session. We sealed the gap between the objective and implant using Blu Tack (Bostik) to prevent any VR projection light from interfering with the GCaMP6f fluorescence signal channel.

### Data analysis (general)

Data analysis was performed using Matlab, unless otherwise noted. Correlation was always the Pearson correlation. Statistical significance was set at 0.05 and was determined using the unpaired t-test, or t-statistic (for testing if a correlation value was different than 0), unless otherwise noted.

### Calcium imaging analysis

Motion correction was done by MOCO (ImageJ plugin, Dubbs et. al. 2016). The reference image for motion correction was selected from the first lap and was used for the rest of the session’s data. Regions of interest (ROIs) were defined in each imaging session in Matlab using custom-written software. Individual neurons were manually selected from a standard deviation image of the first lap (100-160 s of imaging data). ROIs (selected pixel ±4 or ±3 pixels) were determined automatically within a 9 x 9 or 7 x 7 pixel region (to avoid signal contamination from nearby cells) of the manually selected cell. An 81 x 81 or 49 x 49 correlation matrix of the first lap fluorescence signals of the pixels with respect to each other was created (excluding the diagonal). Pixels with low correlation (below 0.45) or high correlation (over 0.85, to avoid non-active regions e.g. cell nucleus) of activity with all other pixels were excluded. If less than 10 pixels remained (in case of cells with few or no calcium transients, i.e. silent cells) then we used the 20 brightest pixels. Fluorescence signals of pixels in each ROI were averaged and used for further analysis. ΔF/F0 was calculated as (F − F0) / F0, where F is the time series of raw fluorescence averaged over the pixels in an ROI, F0 is the mode of a kernel density estimate (Matlab function “ksdensity”) of F within a lap (calculation was done for each lap). ROIs from multiple days were aligned manually using custom-written software in Matlab by matching across the superimposed standard deviation images from pairs of days.

### Detecting calcium activity

Prior to detecting calcium transient events, we applied a Savitzky-Golay filter with 500 ms window (“sgolayflit” function in Matlab) on the ΔF/F0 trace of individual cells. We set the threshold for detecting a calcium event for each neuron at each point in time as the median plus three times the interquartile range of the fluorescence activity of a ±8 s sliding frame (Malvache et. al. 2016). The calcium transient event was defined as the trace from the time of the threshold crossing until the trace fell below the threshold.

### Determination of place fields

First, we rejected activity during periods when the forward running speed of the mice did not exceed 2 cm/s to exclude activity when the animal stopped or was grooming. A place field was defined as a region that had calcium transient events in ≥2 laps whose extents came within 30 cm each of other in the first 4 laps (since there were always at least 4 laps in a given environment in a given session). To fit the distribution of the number of place fields per cell, we fit the Matlab Poisson probability density function (for equal Poisson propensity model) and Negative binomial probability density function (for gamma-Poisson propensity model). The first and last 30 cm of the track were excluded since the animal was locked in (virtual) place at the beginning of each track at the start of each lap for 5 s, and there was an identical end-of-track cue in all environments.

### Determination of reward fields

A place field occurring in a fixed reward area (i.e. the field width overlapped the reward area) was called a reward field. Each reward area included the 25-cm-long reward delivery area, and 25 cm before and 75 cm after the delivery area, i.e. a 125-cm-long segment, which was empirically determined to be the region in which the place field density was increased (Figure 5C).

### Place field number correlation

The place field number correlation was computed as the Pearson correlation coefficient between the place field numbers of the cells in 2 environments/sessions. Silent cells were included in this calculation.

### Population vector correlation

The population vector correlation analysis measured the similarity of a pair of place field representations to assess the degree of stability of the representation in the same environment across sessions and the degree of global remapping between different environments. The binary place field population vectors were determined in each 60 cm spatial bin of each environment (which could also be the same environment in 2 different sessions). The Pearson correlation coefficient was calculated for each pair of corresponding population vectors in the 2 environments and the average across all spatial bins was determined. Silent cells were also included for this analysis. For data from two different sessions, we used only cells which were tracked across both sessions. The first bin and last bin of the track were excluded since the animal was locked in (virtual) place at the beginning of each track at the start of each lap for 5 s, and there was an identical end-of-track cue in all environments.

### Place field shift analysis

We calculated the individual place field shift distance (i.e. the distance that a place field shifted over time within the same environment) by considering every possible pair of place fields of a given cell in 2 sessions. If cell had 2 place fields on one day and 3 place fields on another, then we calculated 6 possible shift distances. The shift distance was measured as the difference between the centers of each pair of place fields. In Figures 2B and 3B, we excluded cell pairs which had a >300 cm shift distance and calculated the proportions in each bin (10 cm) as a function of the time gap between sessions.

### Bayesian decoding

We decoded position of the animal using a standard Bayesian maximum log-likelihood approach (Davidson et al., 2009), in which we assume cells exhibit independent activity, Poisson noise, and a flat prior over the location of the animal.

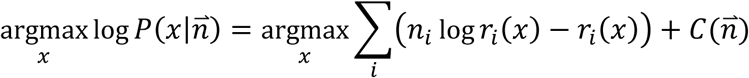

where *x* is the position of the animal, 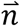 denotes the population vector activity, *i* indexes neuron identity, *C* depends on the population vector activity but does not depend on *x* (thus *C* is irrelevant for the maximum log-likelihood estimation), and *r_i_*_(*x*)_ is a calcium event rate map that was determined for each cell by discretizing the environment into 30 cm segments and averaging a binary variable defined as 1 when the calcium activity was above threshold and 0 otherwise across the times the animal was in that spatial bin. The population vector, 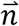, used to decode the activity was the summed activity of the binary neural activity vector over 1s preceding the present position of the animal while its running speed exceeded 5 cm/s, and the rate was scaled to match the size of the time window. The rate map and decoded activity did not necessarily come from the same recording sessions (e.g. using the rate map from an environment on one day to decode the location of the animal in that same environment on a different day). The median decoding error was used as a measure of decoding performance, and chance performance for a decoder that guesses randomly is 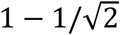 times the environment size by this measure.

### Immunohistochemistry for GABAergic neurons

GABAergic interneurons were stained in two GCaMP6f mice. Mice were perfused transcardially using 4% paraformaldehyde in phosphate buffered saline (PBS). Brains were kept in PBS containing 4% PFA for 24 h at 4 C. and washed with PBS before slicing. 40 µm thick coronal sections were made with a vibratome (Leica, VT1200S) and immersed in PBS. Brain slices were immersed in PBS containing 0.5% Triton X-100 (vol/vol) for 20 min at room temperature and then were transferred to a blocking solution (5% donkey serum (vol/vol), 0.1% Triton X-100 in PBS) for 2 h. Subsequently slices were incubated with primary antibodies in blocking solution for 24 h at 4 C. The primary antibodies were mouse monoclonal anti-GAD67 diluted 1:1000 (MAB5406; Millipore). After incubation, slices were washed three times for 10 min with PBS containing 0.1% Triton X-100 and were incubated with secondary antibodies in the blocking solution for 1 h. The secondary antibodies were Alexa Fluor 647-conjugated anti-mouse IgG2a antibody diluted 1:400 (A-21241; ThermoFisher Scientific). Slices were rinsed three times for 10 min with the PBS containing 0.1% Triton X-100 then were mounted on microscope slides. The fluorescence signal was visualized by the confocal microscopy (Nikon Eclipse Ti) using a CFI Plan Fluor 40X Oil NA 1.3 WD 0.2 objective.

### Place field number, across session model (Figures 4D and 4E)

Place field numbers, *n*, across the population in a given environment during a given imaging session were fit using maximum likelihood to a gamma-Poisson distribution,

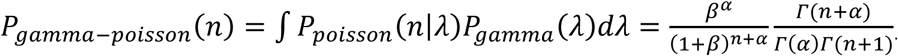

where *Γ* denotes the Gamma function, 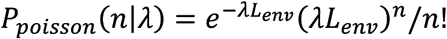 is a Poisson distribution, and the two resulting parameters *α*, *β*, provide a distribution of place field propensities (*λ*) that are gamma distributed,

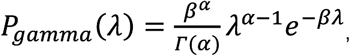

for each environment and session. In our model, we allowed these parameters to be different across sessions, although little variability was seen (Figure S2).

Up to this point, we have described characterizing activity within single sessions, but to characterize cross-session activity, we must connect activity across sessions. The most correlated the activity could be given these constraints would be if the cell propensities had their ordering preserved (i.e. for every pair of neurons *m*, *n* and sessions *s*, *t,* the propensity relationship 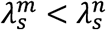 implies that 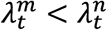). If we consider the cumulative distribution function variables, *X_s_*, *X_t_*, the joint distribution describing propensities for a cell in two different sessions under this order-preserving condition would be *P*(*X_s_*, *X_t_*) = *δ*(*X_s_* − *X_t_*). The full joint distribution of place field numbers for all cells in both environments is then uniquely determined:

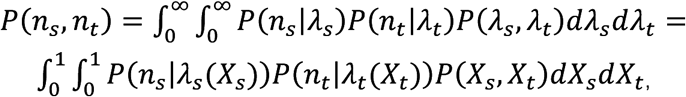

where *λ_s_(X_s_*) is the inverse CDF. Note that the joint distribution in propensity space may be written:

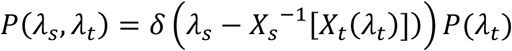

with *X_s_(λ_t_)* denoting the CDF in session *t* and 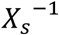 denoting the inverse CDF in session. With this joint distribution, we sampled the predicted number distributions 500 times to measure the correlations and their variability (plotted using 2 standard deviations in each direction) due to neuronal sample size.

### Place field number, reward field number model (Figure 6)

Because we observed significant correlations between reward field numbers and non-reward place field numbers, we sought to determine whether these correlations could be explained by the gamma-distributed propensity that matched the observed place activity. We modeled the non-reward place activity using a gamma-Poisson distribution, and fit a gamma distribution (2 parameters) to the non-reward place field activity observed during the session with no reference to reward activity. Because it is observed that overall population activity increases in reward contexts by a significant amount, we modeled the reward field propensity, *λ_r_*, as a constant gain factor, *G*, multiplying the propensity of the cell in a non-reward place context:

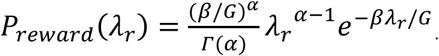

The probability of having no reward field in some region of space declines exponentially for a spatially Poisson process, and so the probability of having a reward field in a single reward area is:

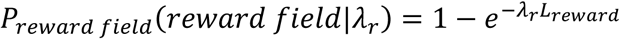

where *L_reward_* is the length of the reward area. If we additionally assume that each reward area has statistically the same properties, then the distribution over reward field number *n_r_*, conditioned on reward propensity will be a binomial distribution:

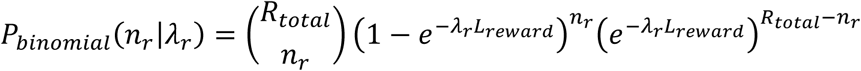

where *R_total_* is the total number of reward areas in the environment. The full distribution describing reward field numbers can be computed, giving rise to a gamma-binomial distribution over reward field numbers:

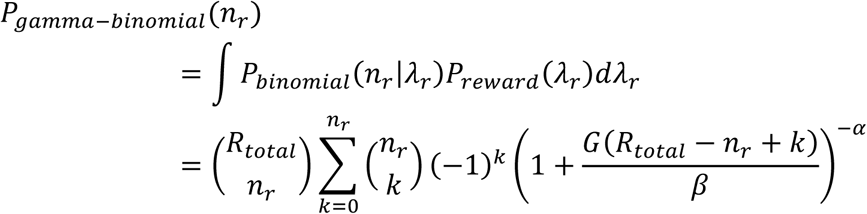

We fit *G* using maximum likelihood to the marginal distribution of reward activity with the gamma parameters fixed to those describing the non-reward place activity. To fully describe the joint distribution of non-reward place and reward activity, the joint distribution must be specified. Given that we observed strong correlations in non-reward place and reward activity, we began by using an analogous order-preserving hypothesis, *P*(*X_p_*, *X_r_*) = *δ*(*X_p_* – *X_r_*), where *X_p_* and *X_r_* are variables representing place and reward propensity, respectively, in CDF space. Using this constant gain model, we observed a very good match for each session for two animals, but two animals had non-reward field – reward field correlations lower than that predicted by the model. To account for these differences, we hypothesized that there may be additional variability, which would result in a broadening of the joint distribution beyond a delta function. To loosen this constraint, we used a copula, a mathematical tool guaranteed to match marginal distributions while varying the underlying joint distribution (Nelsen, Roger B. (1999), *An Introduction to Copulas,* New York: Springer, ISBN 978-0-387-98623-4). A copula defines a multi-dimensional distribution with marginal distributions that are uniform distributions on the interval [0,1]. These uniform marginal distributions may be easily mapped to any marginal distribution using its inverse CDF. The probability density may be written:

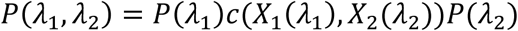

where *P(λ_i_*) describes each marginal distribution, *X_i_*(*λ_i_*) is the CDF evaluated at *λ_i_*, and *c* is the copula density. We chose simple copulas whose single continuous parameter allows the joint distribution to vary from a delta function (perfect order preserving) to a uniform distribution (no correlation between reward and non-reward place activity). After testing the Clayton, Frank, and Gumbel copulas, we observed that the Clayton copula best matched the data (as assessed by the Jensen-Shannon divergence):

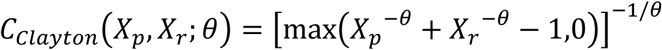

where the copula parameter, *θ*, was chosen to match the correlation observed from the data (after fixing the gain and gamma distribution parameters that define the marginal distributions). In sum, this model contains four parameters: two representing the gamma distribution describing non-reward place activity, one representing the reward-based gain, and one representing the degree of correlation between the reward and non-reward place activity. The model makes predictions about the full joint distribution, which we simulated and compared to the experimentally observed sessions using the Jensen-Shannon divergence:

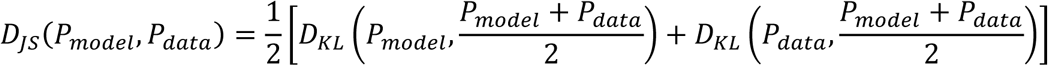

where *D_KL_* denotes the Kullback-Leibler divergence between two models, which is a measure of how distinguishable one probability distribution is from another. The Jensen-Shannon divergence is a measure of how distinct two distributions are from one another, and is preferable to the Kullback-Leibler divergence when one distribution has support over many more outcomes (i.e. a model which predicts small but finite probabilities for unlikely events) than are observed in the other (i.e. data) due to finite sampling. Additionally, we compared the Jensen-Shannon divergence between the observed and modeled results for an alternative model that did not assume a constant gain, but instead allowed the reward propensity to be a nonlinear function of the non-reward field propensity. Specifically, we allowed *λ_r_* = *G*(*λ_p_*)*λ_p_*, where 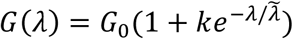, and fit the resulting parameters, *G*_0_, *k*, and 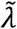, using a maximum-likelihood framework to the full observed joint distribution, *P_data_*(*n_p_*,*n_r_*) after fixing the gamma distribution parameters for non-reward place activity (5 total parameters). We also evaluated a power law distribution for the gain with the same number of parameters, but the exponential form performed better in terms of matched correlations and the Jensen-Shannon divergence. The results showed that the constant gain model with Clayton copula was a better match to the data (measured by the Jensen-Shannon divergence) than this exponential gain model that notably had more parameters and was fit to match the full joint distribution observed in the data (Figure 6).

